# Growth Arrest of Thoracic Aortic Aneurysms in Aging Marfan Mice

**DOI:** 10.1101/2025.06.18.660413

**Authors:** Dar Weiss, Colin Means, Gavin Mays, Nicola Yeung, Cristina Cavinato, Edward P. Manning, TuKiet T. Lam, Jay D. Humphrey

## Abstract

There is a pressing need to identify pathologic mechanisms that render a thoracic aortic aneurysm susceptible to continued enlargement, dissection, or rupture, but additional insight can be gleaned by understanding potential compensatory mechanisms that prevent disease progression and thereby stabilize a lesion. Our biomechanical data suggest that the ascending aorta within a common mouse model of Marfan syndrome, *Fbn1^C1041G/+^*, exhibits progressive disease from 12 weeks to 1 year of age, but near growth arrest from 1 to 2 years of age. Comparison of the biomechanical phenotype, histological characteristics, proteomic signature, and transcriptional profile from 12 weeks to 1 year to 2 years suggests that numerous differentially expressed genes (including downregulated *Ilk*, *Ltbp3,* and *Rictor*) and associated proteins may contribute to late-term growth arrest. There is also a conspicuous absence of proteins associated with inflammation from 1 to 2 years of age. Although there is a need to understand better the interconnected roles of temporal changes in differential gene expression and protein abundance, reducing mTOR signaling and reducing excessive inflammation appears to merit increased attention in preventing continued aneurysmal expansion in Marfan syndrome.

## INTRODUCTION

Marfan syndrome (MFS) is the most common genetic condition that predisposes to thoracic aortopathy, namely, dilatation, dissection, and rupture (Milewicz et al., 2021). MFS results from pathogenic variants in the gene (*FBN1*) that encodes the elastin-associated glycoprotein fibrillin-1. Among other characteristics, increased aortic diameter and stiffness have long been thought to be predictors of lesion progression (Nollen et al., 2004; Selamet et al., 2018). A clinical study of over 200 thoracic aortopathy patients further revealed a critical role of aging both in increasing pulse wave velocity (PWV), the clinical gold standard metric of diffuse stiffening, and decreasing thoracic aorta distensibility (*D*), an indicator of local aortic compliance, in healthy age-matched controls as well as in MFS, bicuspid aortic valve, and familial aortopathy patients (de Wit et al., 2013). Of particular interest, a direct comparison of control vs. MFS data at comparable blood pressures revealed that the Marfan aorta was significantly stiffer than age-matched controls for individuals less than 40 years of age but of comparable stiffness for those greater than 40 years of age, consistent with the suggestion that aortic remodeling in MFS may represent a form of accelerated early aortic aging.

One of the most critical clinical questions is whether a diagnosed thoracic aortic aneurysm (TAA) is expected to progress to dissection or rupture, in which case prophylactic surgery is recommended, or if the lesion is expected either to enlarge slowly without catastrophic failure or to stabilize at a new diameter. Notwithstanding significant insight inferred from clinical data, mouse models of MFS continue to contribute to our understanding of the pathogenesis and natural history of TAAs. The two most common mouse models of MFS are the *Fbn1^mgR/mgR^* model (Pereira et al., 1999), which is hypomorphic for normal fibrillin-1, and the *Fbn1^C1041G/+^* model (Judge et al., 2004), which mimics a human missense variant. The *Fbn1^mgR/mgR^* model presents with an early, severe aortic phenotype while the less-severe *Fbn1^C1041G/+^* model better represents the transcriptional profile seen in human aortas (Sun et al., 2023). It has long been thought that the many differentially expressed genes (DEGs) in the aorta secondary to an underlying pathogenic variant in *FBN1* / *Fbn1* contribute to disease progression, but it is now becoming evident that some gene products resulting from DEGs may serve as protective compensations (Weiss et al., 2023). We thus submit that there is a need not only to determine mechanisms that drive disease progression, but also those that can yield stable growth or growth arrest.

Data from ultrasound, histology, multiphoton microscopy, and ring myography were recently reported for female and male *Fbn1^C1041G/+^* mice at 3, 6, 9, and 12 months of age (Gharree et al., 2022). Aortic diameter increased from 3 to 6 to 9 months of age but appeared to stabilize thereafter; unfortunately, results on possible changes in stiffness were compromised by the methods used. Although it was suggested that “aging exacerbates the disease state especially for males,” effects of aging in mice are expected to manifest well after 1 year of age (Ferruzzi et al., 2018; Rivera et al., 2023). In this study, we quantify transcriptomic, proteomic, histological, and biaxial biomechanical changes in female and male *Fbn1^C1041G/+^*mice from 12 weeks to 1 year to 2 years of age to study possible mechanisms of growth stabilization in the Marfan aorta.

## METHODS

### Animals

All live animal protocols were approved by the Institutional Animal Care and Use Committee of Yale University. Female (F) and male (M) C57BL/6J wild-type (WT) control mice and *Fbn1^C1041G/+^* Marfan syndrome (MFS) mice on a C57BL/6J background were obtained from Jackson Laboratories and inbred locally to maintain colonies for study. Male *Fbn1^C1041G/+^* mice were bred with female C57BL/6J mice to yield heterozygous MFS mice. Consistent with ARRIVE guidelines, mice were randomized to three groups for aging studies (12 weeks, 1 year, or 2 years) as well as three groups for phenotyping (histo-mechanics, proteomics, or transcriptomics). At the scheduled endpoint, tail-cuff blood pressures were measured, then the mice were euthanized via CO_2_ asphyxiation followed by exsanguination upon removal of the thoracic aorta for testing. Histo-mechanical data were collected at {12 weeks, 1 year, 2 years} of age for *n* = {6,6,6} female WT, *n* = {8,5,9} male WT, *n* = {5,10,10} female MFS, and *n* = {5,8,7} male MFS aortas, respectively. Similarly, mRNA was collected from the thoracic aorta from *n* = {4,5,5} female MFS and *n* = {4,2,4} male MFS mice and protein was collected from *n* = {5,5,5} female MFS and *n* = {4,5,5} male MFS mice.

### Biomechanical Testing and Analysis

The active (with smooth muscle cell contraction) and passive (without smooth muscle cell contraction) biomechanical phenotype was characterized using a custom biaxial testing device and validated methods (Ferruzzi et al., 2013; Cavinato et al., 2021). Briefly, the ascending aorta was cleaned of excess perivascular tissue and cannulated with two custom-drawn glass micropipettes – through the aortic root and brachiocephalic artery – and secured with 7-0 suture. The remnant portion of the aortic arch was ligated with sutures between the brachiocephalic and left common carotid artery to enable pressurization. The sample was then placed within the testing chamber and allowed to equilibrate for ∼15 minutes in a heated (37°C) and oxygenated (95/5% O /CO) Krebs-Ringer bicarbonate buffered solution containing 2.5 mM CaCl_2_. The vessels were then subjected to a series of isobaric (luminal pressure of 90 mmHg) - axially isometric (fixed specimen-specific in vivo axial stretch) protocols defined by exposure to 100 mM KCl followed by relaxation (KCl washed out), 1 μM AngII followed by relaxation (AngII washed out), 1 μM phenylephrine (PE), then without washout 10 µM acetylcholine (Ach – an endothelial-dependent source of nitric oxide) and finally 1 mM N_ω_-Nitro-L-arginine methyl ester (L-NAME -- a blocker of endogenous production of nitric oxide).

Next, the Krebs solution was replaced with a Hanks buffered salt solution (HBSS) at room temperature to ensure a passive response. Vessels were preconditioned via four cycles of pressurization between 10 and 140 mmHg while held fixed at their individual in vivo axial stretch. They were then subjected to three cyclic pressure-diameter (*P-d*) protocols, with luminal pressure cycled between 10 and 140 mmHg while axial stretch was maintained fixed at the in vivo value, then ±5% of this value, and four cyclic axial force-length (*f-l*) tests, with force cycled between 0 and a force equal to the maximum value measured during the pressurization test at 5% above the in vivo axial stretch, while the luminal pressure was maintained fixed at 10, 60, 100, then 140 mmHg. Pressure, axial force, outer diameter, and axial length were recorded on-line for all seven of these protocols and the last unloading curve of each was used for subsequent data analysis (∼2800 data points per sample).

Consistent with prior studies on MFS aortas (Cavinato et al., 2021; Weiss et al., 2023), we employed a radially homogenized “four-fiber family” form of a strain energy function w (having unit of kPa) to characterize the passive biaxial properties of the aortic wall, with

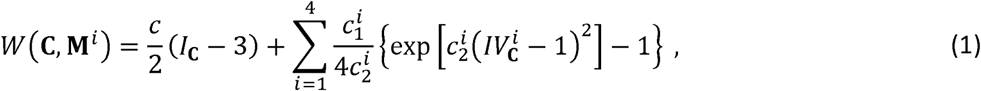

where *c* (kPa), *c^i^*_1_ (kPa), and *c^i^*_2_ (-) are material parameters and superscript ; denotes four families of collagen fibers having predominantly axial (*i* = 1), circumferential (*i* = 2), and symmetric diagonal (*i* = 3,4) orientations. **C** = **F^T^F** is the right Cauchy-Green tensor, where **F** = diag(λ_*r*_, λ_*θ*_, θ_*z*_) is the experimentally measured deformation gradient tensor with tissue incompressibility requiring λ_*r*_ = 1/(λ_θ_λ_*Z*_). **M^*i*^ = [0, sin α^*i*^_0_, cos α^*i*^_0_**] is a unit vector in the direction of the *i*th fiber family, with fiber angle ai defined relative to the axial direction in a reference configuration. Thus, axial and circumferential families are oriented at α^1^_0_ = 0° and α^2^_0_ = 90°, respectively, while the symmetric diagonal fiber orientation a3,4 is determined from data. Best-fit values of the 8 parameters are listed in Supplemental **Table S1**. Because blood pressures did not differ significantly between WT and MFS mice, all mechanical metrics were compared at specimen-specific values of the in vivo axial stretch but the same distending pressures (80 – diastolic, 100, and 120 – systolic values in mmHg). Clinical distensibility was computed as *D* = (*d_sys_* - *d_dias_*)/(*P_sys_* - *P_dias_*)*d_dias_* where *d* and *P* are inner diameter and pressure, respectively.

### Multiphoton Microscopy

Function derives from structure. Following mechanical testing, vessels were immersed in HBSS at room temperature and held at the specimen-specific axial stretch and a common intraluminal pressure of 80 mmHg. A two-photon (LaVision Biotec TriMScope) microscope with a TiSa laser tuned at 820 nm was used with an Olympus 20X objective lens (N.A. 0.95). Three signals were second harmonic generation for fibrillar collagen (390-425 nm), two-photon excited fluorescence for elastin (500-550 nm), and SYTO17 red fluorescence of cell nuclei (above 550 nm). Three-dimensional images were acquired with an axial-circumferential field of view of 500 μm × 500 μm at a consistent location corresponding to the center of the outer curvature, with a numerical imaging resolution of 0.48 μm/pixel and out-of-plane (radial axis) step size of 1 μm/pixel. All images were processed as described previously (Cavinato et al., 2021). A three-dimensional elastin porosity was defined as the ratio of the volume identified as voids and the volume occupied by elastic fibers – including lamellar and intra-lamellar. Description of collagen fiber bundles focused on in-plane (axial-circumferential) parameters of fiber straightness and bundle width.

### Histology

Following mechanical testing and multiphoton imaging, each vessel was fixed overnight in a 10% neutral buffered formalin solution, then stored in 70% ethanol at 4°C until embedding in paraffin and serial sectioning at 5 µm. Mounted slides were stained with either Movat’s pentachrome (MOV), to detect elastin (black), collagen (yellow/brown), cytoplasm (red/purple), mucoid material (glycosaminoglycans - aqua/blue), and fibrin (pink/red), or picro-sirius red (PSR), to detect fibrillar collagens in a spectrum from thick (red/orange) to thin (yellow/green). MOV sections were used to compute medial and adventitial wall areas as well as areas for elastin, cytoplasm, collagen, glycosaminoglycans, and fibrin (Cavinato et al., 2021; Chen et al., 2023).

### Proteomics

Having characterized differences in histo-mechanical characteristics between age- and sex-matched WT and MFS mice, we next focused on possible changes in proteomic signature in the MFS aorta as a function of age. We first used 300 μL of a stock radioimmunoprecipitation assay buffer, containing protease (Pierce #87786 at 1% of lysis buffer) and phosphatase inhibitor (Pierce #78420 at 2.5% of lysis buffer) cocktails, to liberate cellular proteins within the aortic wall. The mixture was ultrasonicated with four 1-sec bursts of 10 amp, then allowed to incubate at 95°C for 10 minutes with vortex every 2 minutes. The mixture was then cooled to room temperature and cysteines were reduced with dithiothreitol (DTT) at 60°C for 30 min, then cooled again to room temperature and alkylated with iodoacetamide (IAN) at room temperature in the dark for 30 min. Proteins were precipitated with acetone. Briefly, 1440 µL of cold -20°C acetone was added to the protein solution and vortexed, then placed at -20°C overnight. The protein pellet (after removal of supernatant post-centrifugation at 16000 g for 10 minutes at 4°C) was washed once with 1 mL of cold -20°C acetone, with repeated centrifugation at 16,000g for 10 minutes at 4°C. The supernatant was discarded, and the protein pellet was air dried for 5 minutes, brought up in 8 M Urea/400 mM ammonium bicarbonate, and digested with PNGase F at 37°C for 5 hours on a shaker. LysC (2 µL at 0.5µg/µL) was added for overnight digestion at 37°C with shaker; trypsin (µL of 0.5 µg/µL) was added and the mixture was incubated for 6.5 hours on a shaker. Digestion was quenched with 20% trifluoroacetic acid (TFA). Samples were desalted using a BioPureSPIN PROTO 300 C18 macro-spin columns (The Nest Group #HMM S18V) and dried via SpeedVac (Thermo Scientific SAVANT RVT-4104). Peptide concentrations were determined using a NanoDrop spectrophotometer (Thermo Scientific NanoDrop 2000) to load 0.3 μg / 5 μL of each sample per column for the liquid chromatography – mass spectrometry (LC MS/MS) analysis.

Label-free quantification (LFQ) / Data Dependent Acquisition (DDA) was performed on a Thermo Scientific Orbitrap Fusion mass spectrometer connected to a Waters nanoACQUITY UPLC system equipped with a Waters Symmetry® C18 180 μm × 20 mm trap column and a 1.7-μm, 75 μm × 250 mm nanoACQUITY UPLC column (35 °C). To ensure a high level of identification and quantitation integrity, a resolution of 120,000 and 60,000 was utilized for MS and MS/MS data collection, respectively. MS (scan range between m/z 350 to 1550 with AGC target of 400,000) and MS/MS (using isolation window of 1.6, and AGC Target of 50,000 with Higher-energy C-Trap Dissociation (HCD) at 28% collisional energy) spectra were acquired using a three-second cycle time with a 30 second Dynamic Exclusion duration. All MS (Profile) and MS/MS (centroid) peaks were detected in the Orbitrap and ion trap regions, respectively. Peptides were in trapping mode on the LC for 3 min at 5 μL/min in 99 % Buffer A (0.1% FA in water) and 1 % Buffer B (0.075% FA in acetonitrile – ACN) prior to eluting with linear gradients that reach 5% B at 5 min, 20% B at 125 min, 35% B at 170 min, and 97% B at 175 min for 5 minutes; then back down to 3% at 182 min. Four blank injections (1^st^ 100 % ACN, 2^nd^ and 3^rd^ are 50:50 ACN:Water, and 4^th^ is Buffer A) followed each sample injection to ensure there was no sample carry over.

A label-free quantitation using Progenesis QI (Nonlinear Dynamics, LLC, UK; version 4.2), similar to that found in Bordner et al. (2011), was used to obtain quantitative information on peptides and proteins. Here, the LC–MS/MS data (uninterpreted MS/MS spectra) were processed with protein identification carried out using an in-house Mascot search engine (version 2.8). The Progenesis QI software performs chromatographic/spectral alignment (wherein one run is chosen as a reference to which all other data files are aligned), mass spectral peak picking and filtering, and quantitation of proteins and peptides. A normalization factor for each run was calculated to account for differences in sample load between injections as well as differences in ionization. The experimental design was set up to group multiple injections (technical and biological replicates from each run) into each comparison sets. The algorithm then calculated the tabulated raw and normalized abundances and ANOVA *p*-values for each feature in the data set. The MS and MS/MS spectra were exported in .mgf (Mascot generic files) format for database searching. The Mascot search algorithm (Hirosawa et al., 1993) was used for searching against the Swiss Protein database with taxonomy restricted to *Mus musculus*. Carbamidomethyl (Cys), oxidation of Met, Phospho (Ser, Thr, Tyr), Deamidation (Asn, Asp), Acetyl (Lys), and Acetyl (Protein N-Term) were entered as variable modifications. Two missed tryptic cleavages were allowed, precursor mass tolerance was set to 10 ppm, and fragment mass tolerance was set to 0.02 Da. The significance threshold was set based on a False Discovery Rate (FDR) of 1%. The Mascot search results were exported as .xml files and then imported into the processed dataset in Progenesis QI software where identified peptides were synced with the corresponding quantified spectral features. Protein abundance was then calculated from the sum of all unique normalized non-conflicting peptide ions for a specific protein on each run.

### Transcriptomics

Finally, we quantified the transcriptional profile in the MFS aorta as a function of age using bulk RNA-seq following our published methods (Spronck et al., 2023). Briefly, RNA was isolated from the thoracic aortas of MFS mice at all three ages using an RNeasyMini Kit (Qiagen) according to manufacturer’s specifications, and whole-transcriptome sequencing was performed by The Yale Center for Genome Analysis using a NovaSeq 6000 System (Illumina, Inc.). Sequenced reads were imported into CLC Genomics Workbench V23 (Qiagen) and, following quality control, reads were trimmed and aligned / mapped to a *Mus musculus* reference genome. Reads were then automatically processed using trimmed mean of M values (TMM) normalization, log counts per million (CPM) transformation, and Z-Score normalization to generate principal component analysis (PCA) and heat map plots (Euclidean distance and complete linkage for clustering). To compare between age groups, expression values were normalized by applying TMM normalization to CPM values – CPM (TMM-adjusted). Genes were then filtered using Automatic Independent Filtering (DESeq2) prior to false discovery rate correction to improve power. Benjamini–Hochberg correction, as implemented in the CLC Genomic Workbench, was used to obtain *p*-values adjusted for multiple comparisons, and genes with an adjusted *p*-value less than 0.05 were defined as differentially expressed genes (DEGs). DEGs were then exported to the QIAGEN Ingenuity Pathway Analysis (IPA) software for gene ontology studies. Comparative analyses were performed to reveal the most significant processes and functions across the different times and to generate corresponding bubble charts and bar plots.

### Statistics

Due to deviations from normality (data failed to pass Kolmogorov-Smirnov normality test), we used the nonparametric Kruskal Wallis one-way ANOVA on Ranks followed by Dunn’s post-hoc test for multiple comparisons to interpret geometric, histo-mechanical and multiphoton microscopy metrics across the 12 study groups. For all reported comparisons, a value of *p*<0.05 was considered significant.

## RESULTS

### Sex as a biological variable

Characteristic vessel-level trends and differences were generally comparable for the ascending aorta from female and male MFS mice relative to sex- and age-matched WT controls, though dilatation and both material and structural stiffening were more severe in male than female MFS mice. For this reason, and to simplify the presentation across 12 groups, comparisons by sex are provided in Supplemental Materials (Supplemental **Figures S1-S3**; Supplemental **Tables S1-S4**) while we focus here on age-dependent differences between male WT and male MFS mice.

### Biomechanical Phenotype

Natural aging of male WT mice to 2 years of age resulted in expected changes in the thoracic aorta phenotype (Rivera et al., 2024): progressive decreases in both elastic energy storage w and distensibility D as well as a decrease in the in vivo value of axial stretch,^iv^ and an increase in the circumferential material stiffness r_ee_ (**Figure 1; Tables S2-S4**). Pressure-diameter data suggested that the male MFS aorta was dilated relative to the age-matched male WT aorta at 12 weeks and especially at 1 and 2 years of age, with percent dilatation similar at the latter two ages (**Figure 1A**). Of the different biomechanical metrics, marked increases in circumferential material stiffness best associate with TAA propensity and progression (Bellini et al., 2017). At 100 mmHg pressure, this stiffness was ∼2.0-fold higher in male MFS mice at 12 weeks of age relative to age-matched male WT mice and ∼1.68-fold higher at 1 and 2 years (**Figure 1B**; **Table S3**). The early increase in stiffness in MFS thus preceded marked aortic dilatation while the latter reduction in fold-increase relative to WT reflected the modest stiffening due to natural aging in WT. Values of elastic energy storage and distensibility were lower in the MFS aorta relative to WT, with values in MFS similar at 1 and 2 years (**Figure 1C,D**). Additional comparisons (**Figure S1; Tables S2-S4**) further suggest that the overall aortic phenotype worsened in the MFS aorta from 12 weeks to 1 year of age, after which age-related changes were more modest.

**Figure 1.**
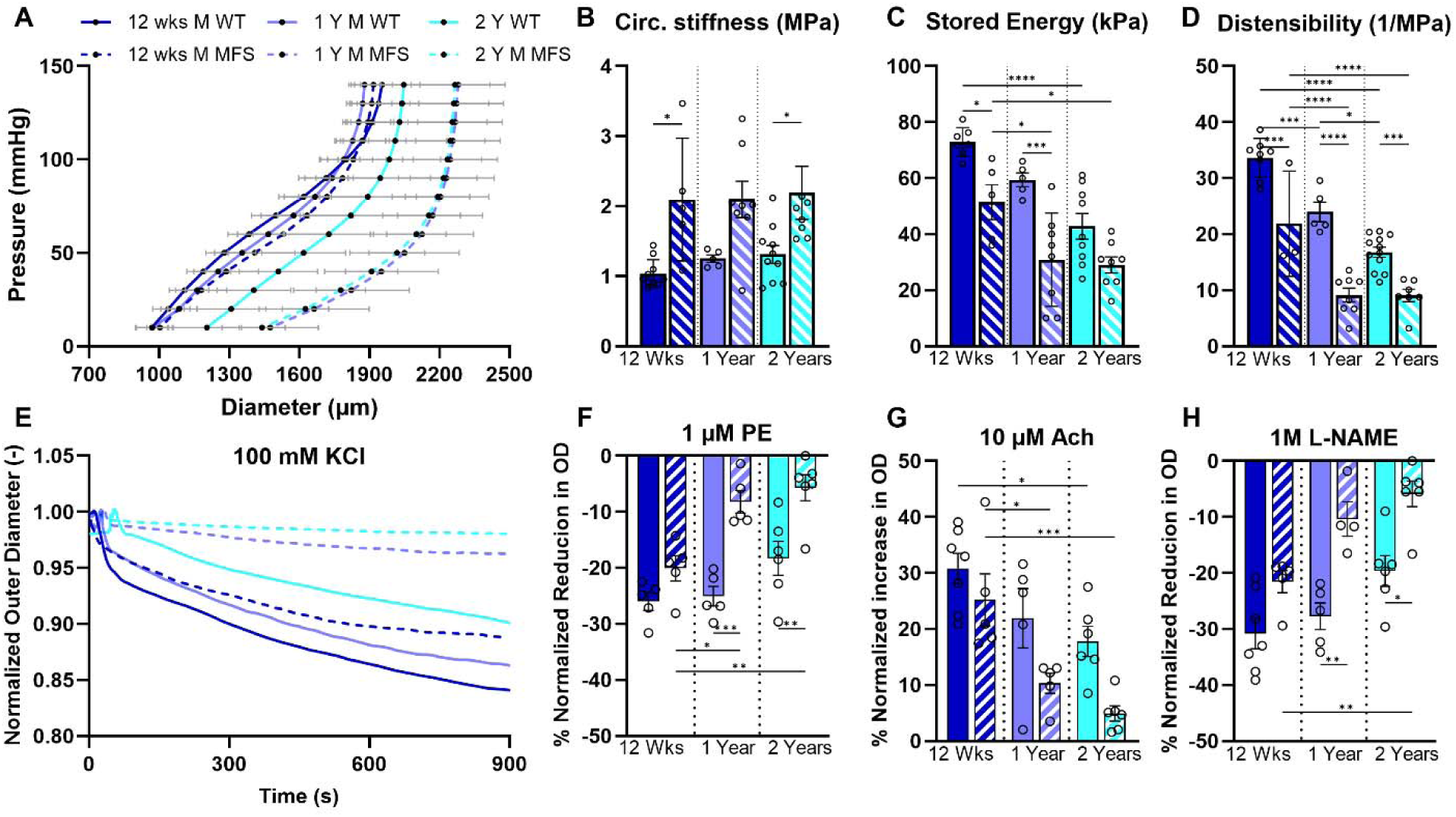
Biomechanical data for the ascending aorta from age-matched male WT (solid lines and bars) and *Fbn1^C1041G/+^* MFS (dashed lines, cross-hatched bars) mice. A. Pressure-outer diameter responses at specimen-specific in vivo stretch. B-D. Circumferential material stiffness, elastic energy storage, and overall distensibility. E. Time-dependent reduction in outer diameter in response to 100 mM KCl. F-H. Diameter reduction after 15 minutes of vasoactive stimulus with phenylephrine (PE), acetylcholine (Ach), and the endothelial-derived nitric oxide synthase-inhibitor L-NAME. Mean ± SEM, with *, **, ***, **** indicating significance at *p* < 0.05, 0.01, 0.001, 0.0001, respectively. See also Supplemental Figures S1 and S2 which contrast females and males. Calculated values are based on material parameters listed in Supplemental Table S1 with further biomechanical metrics listed in Tables S2-S4.

Natural aging also resulted in modest decreases in vasoactive capability whereas MFS resulted in greater decreases when compared to age-matched WT controls (**Figures 1E, S2**). Focusing on vasoconstriction to 10 μM PE, vasodilation to 10 μM Ach, and vasoconstriction to 1mM L-NAME (**Figure 1F-H**) revealed that the diminished function in MFS at 1 year of age persisted rather than worsened at 2 years of age. Taken together, the passive and active biaxial data suggested that the aortic phenotype worsens from 12 weeks to 1 year of age in *Fbn1^C1041G/+^* MFS mice but thereafter tends to remain relatively stable as the mice continue to age to 2 years old.

### Aortic Composition

Standard histological assessments of MOV- and PSR-stained cross-sections (**Figures 2, S3**) revealed that natural aging of the male WT thoracic aorta resulted in modest changes, notably small differences in GAGs but slight increases in mural cytoplasm and fibrillar collagens (**Figure 3B-D**). By contrast, mural GAGS were higher in MFS relative to WT at 1 and 2 years of age. Increased GAGs can play diverse roles in aortic mechanics and mechanobiology (Roccabianca et al., 2014). Age-matched MFS aortas also showed higher mural cytoplasm and higher fibrillar collagens than WT controls; importantly, these increases mainly occurred from 12 weeks to 1 year while showing little change thereafter to 2 years (**Figure 3B-D**). These findings were consistent with both the greater wall thickness and higher material stiffness in MFS relative to WT at 1 and 2 years of age.

**Figure 2.**
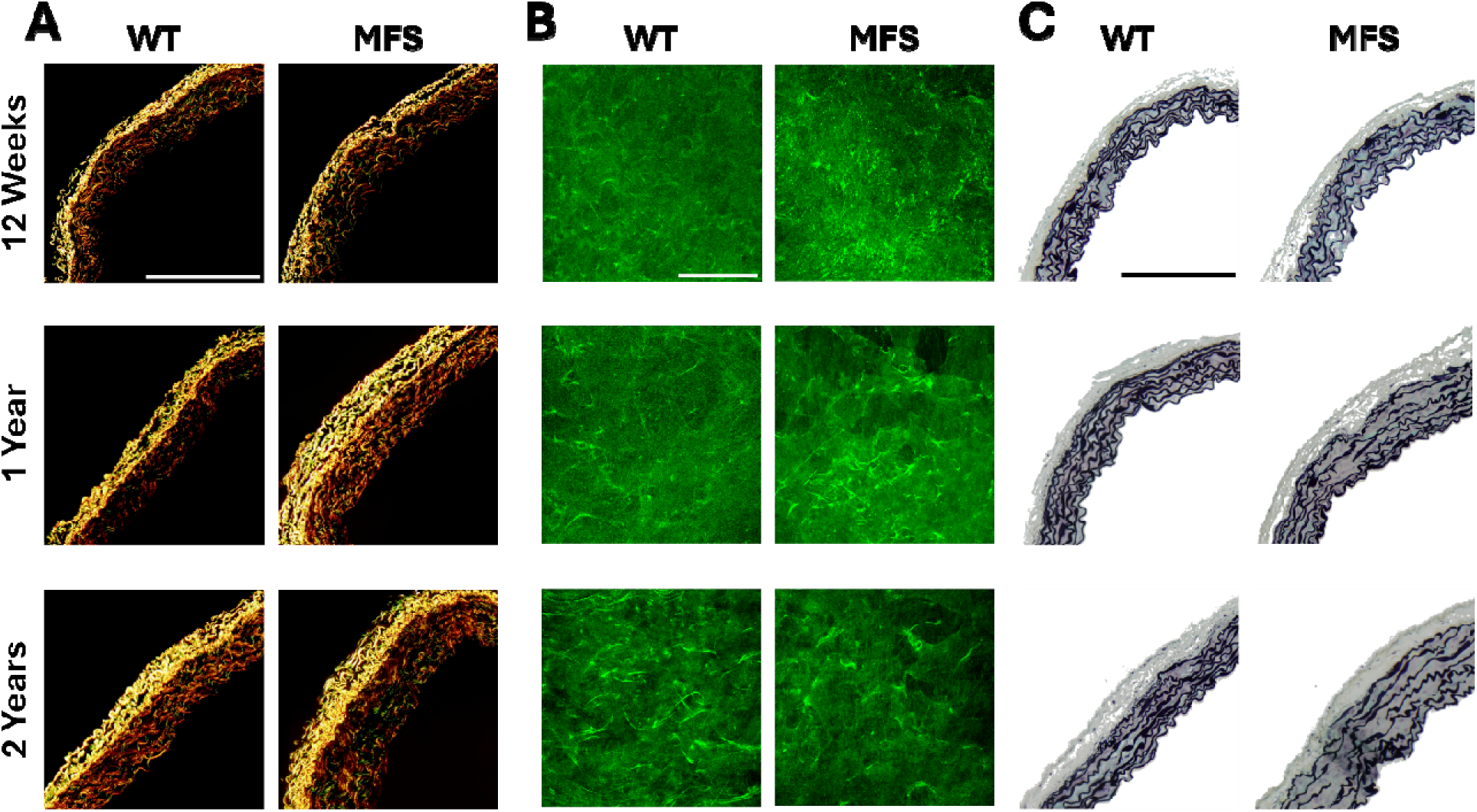
Representative histological images for male WT and MFS aortas at all three ages. A. Polarized light microscopic images of collagen (red/orange = thick fibers, yellow/green = thin fibers) from picrosirius red stained sections. B. Two-photon fluorescence images of elastin from multiphoton microscopy. C. Bright field light microscopic images of Movat-stained sections: elastin (black), collagen (grey/yellow), glycosaminoglycans (blue), and cytoplasm (pink/red). See also Supplemental Figure S3. Scale bar = 100 µm.

**Figure 3.**
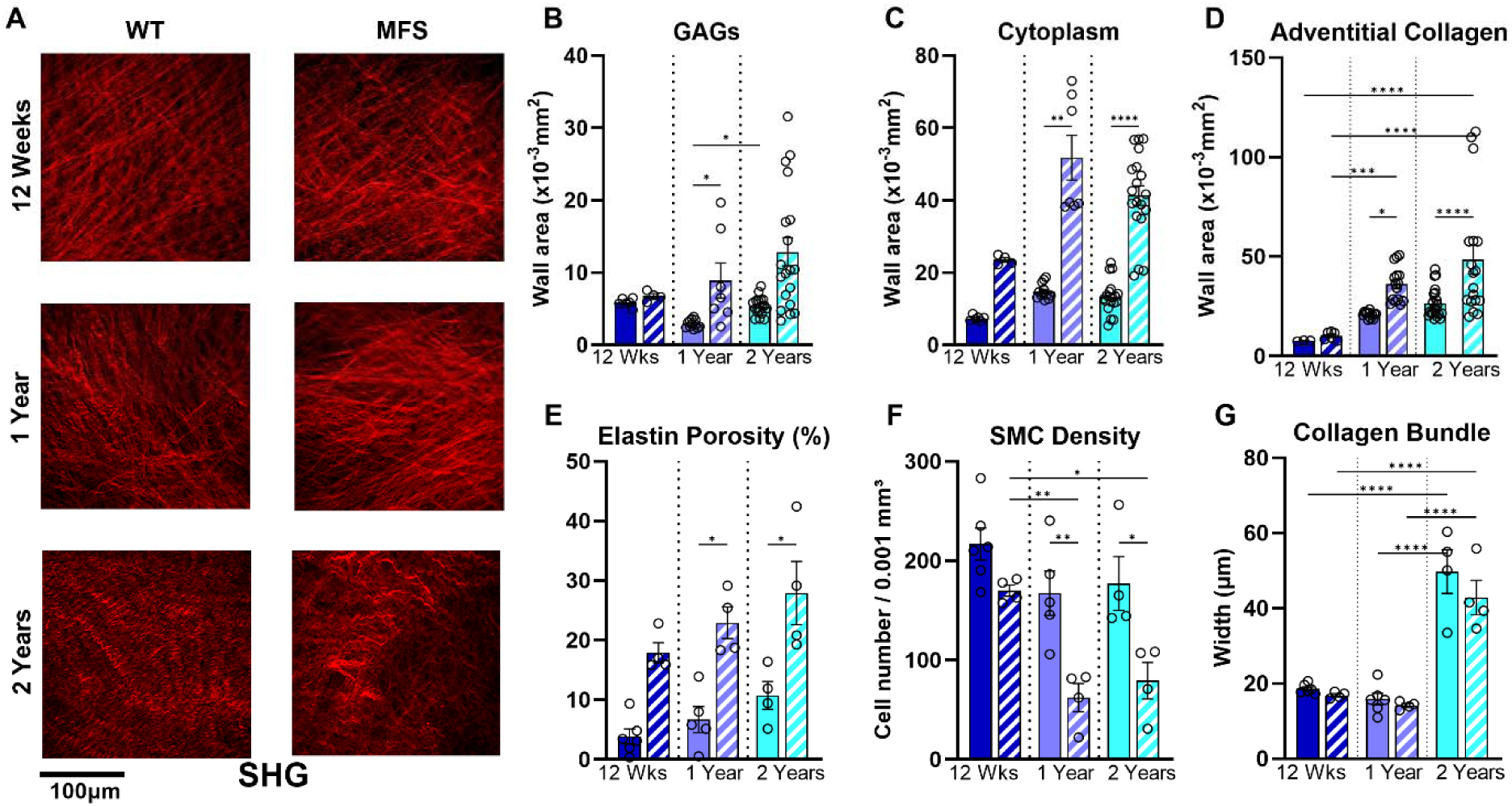
Additional histological / microstructural information for male WT (solid) and MFS (cross-hatched) aortas at all three ages, including (A) second harmonic generation of fibrillar collagen from multiphoton imaging and (B-G) quantification of select constituents.

Multiphoton microscopy revealed further that natural aging resulted in modest progressive increases in elastin porosity whereas MFS exacerbated these increases at each age (**Figures 2, 3E**). Smooth muscle cell density (based on nuclei) decreased slightly with natural aging but was much lower in MFS than in age-matched controls, with the reduced values in MFS similar at 1 and 2 years of age (**Figure 3F**). These findings are consistent with the diminished vasoactive findings of **Figure 1** that could also stem from a transition from a contractile to synthetic smooth muscle phenotype. Finally, second harmonic generation (**Figure 3A,D**) revealed that collagen fiber bundle width was greater at 2 years of age than at 12 weeks and 1 year of age for both WT and MFS, with little difference by genotype (**Figure 3G**). Albeit not shown, collagen fiber alignment changed little with age in both WT and MFS aortas. Overall, most changes in the Marfan aorta again occurred between 12 weeks and 1 year of age.

### Proteomic Signature

Alterations in mural composition should be reflected by an altered proteome. Given the provocative finding that multiple histo-mechanical changes in the male MFS aorta tended to progress from 12 weeks to 1 year of age and then stabilize, we focus here on the evolving MFS proteome. Principal component analysis (PCA) revealed some separation in differential protein abundance (DPA) in MFS at each of the three ages while volcano plots revealed greater differences between 1 and 2 years than between either 1 year and 12 weeks or 2 years and 12 weeks (Supplemental **Figure S4a-e**; **Table S5**). DPA at 1 year relative to 12 weeks included, among others, statistically significant changes in (**Figure S4e**; alphabetically) ACAN, ITGA1, ITGAV, ITGB3, LOX (decreased), LOXL3 (largest increase of proteins considered), PDGFRA, PPP1R12A (decreased), PRKG1 (decreased), TIMP3, and VCAN; further differences at 2 years relative to 1 year included BGN, COL5A1 (decreased), COL12A1 (largest increase), COL18A1, FN1, ITGAM, ITGA5, ITGB5, LGALS3, LTBP3 (decreased), MMP2, PXN, SPARC, THBS1, TNFRSF11B, TNS1, and VCAN. Many of these proteins contribute to the extracellular matrix and its mechanosensing by the local cells. Temporal changes in 60 select proteins across multiple classes – contractile, extracellular matrix (ECM), focal adhesion, matricellular, proteolytic, and immune – found to be potentially important in diverse studies of mouse models of aortopathy are highlighted in **Figures 4 and S5a-c**. Whereas the reader is encouraged to peruse these data, first consider a few proteins motivated by specific histo-mechanical findings. Notwithstanding progressive increases in fibulin-5 and fibrillin-1 despite near constant elastin in the proteome, the progressive increase in MMP-2 may have reflected the multiphoton-detected progressive increase in elastin porosity, noting however that TIMP-3 similarly increased. Despite a near constant fibrillar collagen III but decreased collagen V, progressive increases in collagen I, biglycan, and lysyl oxidase like-1 appeared to reflect early increases observed histologically in medial and especially adventitial collagen, which in turn appeared to reflect increases in circumferential material stiffness. Remodeling of adventitial collagen can be compensatory in TAAs (Kawamura et al., 2021); whereas smooth muscle cells are generally assumed to be responsible for matrix changes in the media, multiple cell types can contribute to adventitial remodeling (Wu et al., 2016). The progressive increase in the aggregating proteoglycans versican and aggrecan also appeared to reflect the diffuse increases observed histologically in the MOV-stained sections. Changes in contractile proteins were less clear. Smooth muscle alpha actin (SMαA), smooth muscle myosin heavy chain (SM-MHC), and myosin light chain 9 changed little while MYPT1 and protein kinase cGMP-dependent 1 (PRKG1), which inhibit contraction, decreased with age; recall from the functional tests that vasoconstriction was yet decreased in the Marfan aorta with aging, which may have reflected loss of cell numbers more than cell function.

**Figure 4.**
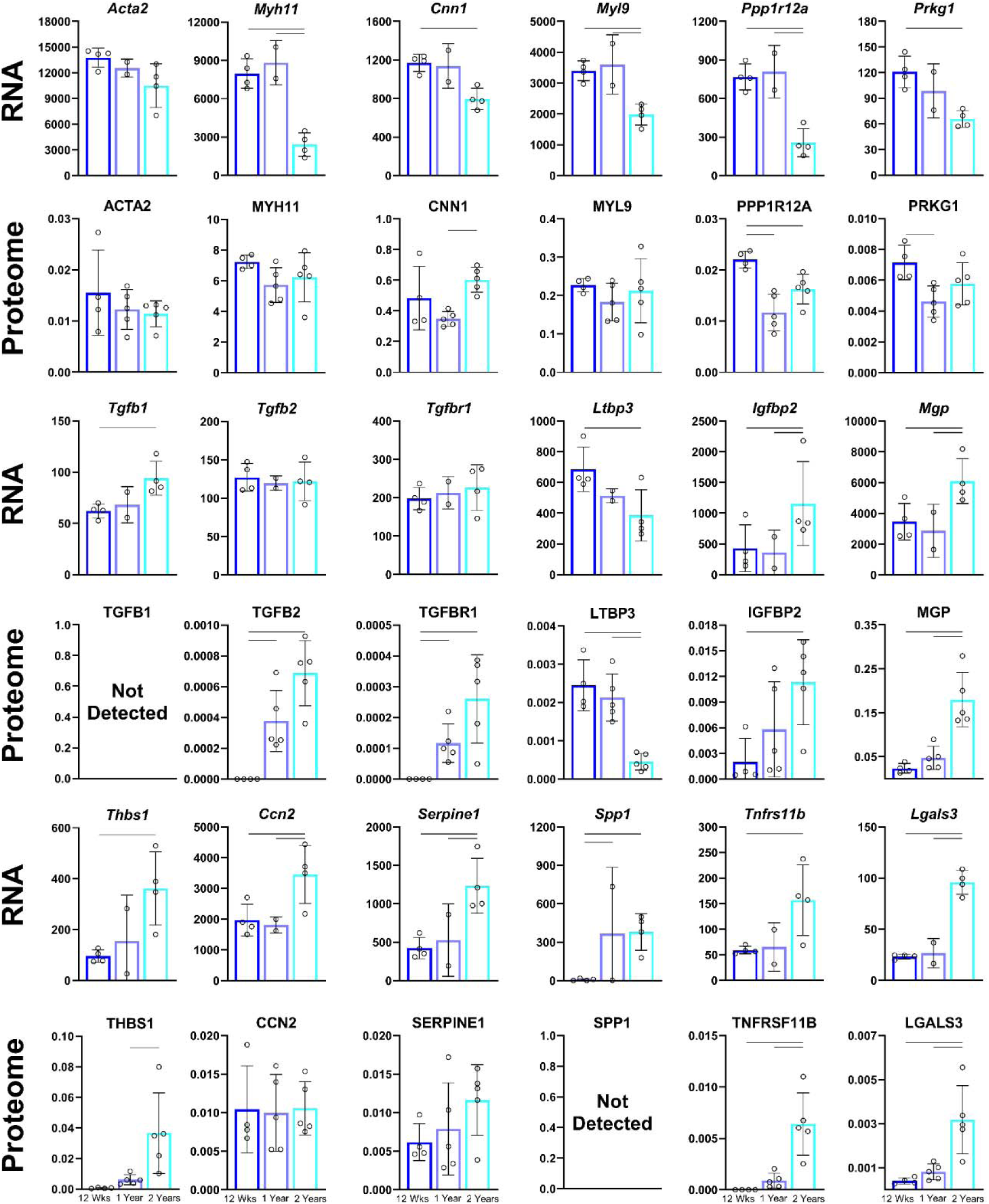
Paired temporal changes in select gene expression (RNA; units of CPM) and protein abundance (proteome) for aortas from male MFS mice at all three ages. See also Supplemental Figure S5A-C for additional constituents, 60 total, and Figure S7 for additional quantification. In each panel, 12 weeks (left bar), 1 year (middle bar), 2 years (right bar). Significance overbars bars indicate *p*<0.05. See supplementary Tables S5 and S6 for detailed *p* values of all comparisons.

Transforming growth factor-beta (TGFβ) signaling affects many gene products, including contractile proteins and matrix synthesis. Both TGFβ2 (ligand) and TGβRI (receptor) increased with age in MFS while latent transforming growth factor binding protein 3 (LTBP3) decreased at 2 years. Absent LTBP3 improves the aortic phenotype in the *Fbn1^mgR/mgR^* mouse model of severe MFS (Zilberberg et al., 2015; Korneva et al., 2019). By contrast, insulin-like growth factor binding protein 2 (IGFBP2) increased consistent with many reports of increased *Igfbp2* in many MFS studies, including the present.

Noting that dysfunctional mechanosensing or mechanoregulation of ECM appear to contribute to disease progression in MFS (Humphrey et al., 2015), many focal adhesion-associated proteins had altered abundance. These include integrin subunits α1, α5, α8, αv, β1, β3, and β5 as well as αm (immune cell). There were associated changes in focal adhesion associated proteins such as paxillin and tensin, but not in talin, vinculin, or filamin-A, and no changes in integrin linked kinase or focal adhesion kinase. Increased matricellular protein thrombospondin-1 (THBS-1) has been implicated in dysfunctional mechanosensing in thoracic aortopathy (Yamashiro et al., 2018); THBS-1 increased with age in MFS as did plasminogen activator inhibitor 1 (PAI-1), which has been suggested to allow aneurysm enlargement but to protect against dissection and thus be “self-protective” (Gomez et al., 2013).

### Transcriptional Profile

Evidence of DEGs by RNA-seq provides insight into what the cell is currently sensing and how it is responding. To complement the proteomics, we similarly focused on age-dependent changes in the MFS transcriptome (**Figure 6**). Expectedly (Gerstein at al., 2003), age-dependent changes in only 33% of 106 selected transcripts – contractile, structural ECM, matricellular, focal adhesion, proteolytic, and immune – correlated positively with changes detected in the proteome (**Figure S7**). Notwithstanding many DEGs seen in both female (**Figure S8**) and male (**Figure S9**) MFS mice across the three ages, similar to many prior studies in both *Fbn1^mgR/mgR^* and *Fbn1^C1041G/+^* models (Bhusan et al., 2018; Chen et al., 2021), we focused primarily on select transcripts in males that are critical in dictating aortic development and integrity (Kelleher et al., 2004; Weiss et al. 2024). Principal component analysis (PCA) revealed some separation in DEGs in MFS across the three ages while volcano plots revealed greater differences between 1 and 2 years than between either 1 year and 12 weeks or 2 years and 12 weeks (**Figures 5, S9; Table S6**). Differences at 1 year relative to 12 weeks yet included, among others, statistically significant changes in (alphabetically) *Ccl2, Ccr2, Col15a1* (downregulated)*, Cx3Cr1, Eln* (downregulated)*, Itgam, Lum, Mmp13, and Spp1*; differences at 2 years relative to 1 year included those for *Ccl2, Ccl5, Cx3Cr1, Cxcl13, Itgam, Lgals3, Myh11* (downregulated)*, Ppp1r12a* (downregulated)*, Rictor* (downregulated), *Serpine1*, and *Tnfrsf11b*. Many of these transcripts appear to suggest altered immune cell activity as well as altered mechanosensing. Temporal changes in 60 select transcripts across multiple classes – contractile, extracellular matrix (ECM), focal adhesion, matricellular, proteolytic, and immune – are contrasted against those for the proteome in **Figures 4 and S5a-c**.

**Figure 5.**
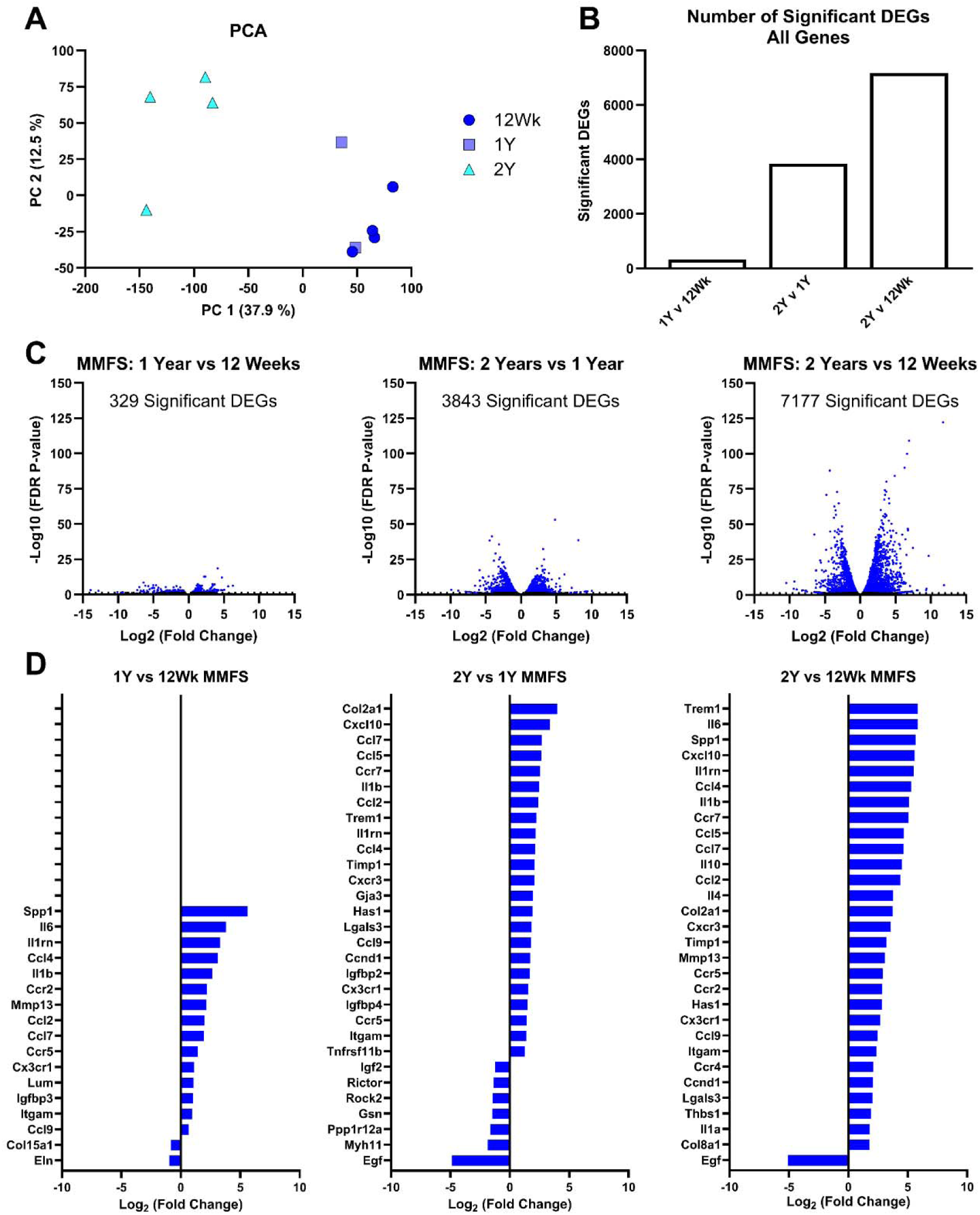
Further quantification of select transcriptomic changes as a function of age in male MFS mice. A. Principal component analysis (PCA). B. Overall number of differentially expressed genes (DEGs) when comparing the Marfan aorta at 1 year vs. 12 weeks, 2 years vs. 1 year, and 2 years vs. 12 weeks. C. Volcano plots show relative up- and down-regulated DEGs for the same three comparisons. D. Log_2_ fold-changes in 30 genes selected based on prior results reported for the Marfan aorta. See also Figure S9.

## DISCUSSION

Fibrillin-1 has diverse roles within the aorta. Among other functions, it contributes to the long-term structural stability of the elastic lamellar structures, the intra-lamellar connections that facilitate mechanosensing, and sequestration of latent TGF-β via its interactions with latent transforming growth factor binding proteins (LTBPs). Most prior studies of the aorta in MFS mice appropriately sought to identify mechanisms that drive disease progression (e.g., Cook et al., 2015; Deleeuw et al., 2021; Chen et al., 2023; Li et al., 2023). Notwithstanding the critical importance of such studies, there is also much to learn from lesions that either grow slowly or not at all. Although focused on abdominal aortic aneurysms, computational (Wilson et al., 2013) and theoretical (Cyron et al., 2014) studies suggest that progressive (unstable) versus arrested (stable) aneurysm growth are distinguished, in part, by mechanobiologically driven changes in aortic composition and material properties. The milder phenotype of the common carotid artery (neural-crest-derived smooth muscle cells) and descending thoracic aorta (mesoderm-derived smooth muscle cells) in MFS mice (Eberth et al., 2009; Westenberg et al., 2011; Schwill et al., 2013; Bellini et al., 2016) may hold some clues regarding slow or stable growth, but we focused on the most affected segment of the aorta (ascending) that yet exhibits a mild long-term phenotype in *Fbn1^C1041G/+^* mice. To study possible growth arrest during aging, we followed these mice for 2 years.

Previously reported histo-mechanical characteristics of the thoracic aorta in aged (≥ 20 months) WT mice include a thickened wall with increased fibrillar collagens and diffusely increased GAGs but decreased cytoplasm, decreased elastic energy storage, decreased in vivo axial stretch, decreased axial stiffness, and decreased distensibility with increased circumferential material stiffness (Ferruzzi et al., 2018; Rivera et al., 2024). Our findings in aging male WT mice were largely consistent with these studies. Prior reports of associated transcriptional changes in aging (≥ 20 months) of WT mice include increased *Col1a1*, *Col3a1*, and *Col6a1* and decreased *Bgn*, *Dcn, Eln*, and *Timp2,3* (Rammos et al., 2014; Rivera et al., 2024), with scRNA-seq further revealing increases in CD3+ T-cells and CD31+ endothelial cells but decreases in smooth muscle cells with age (Rivera et al., 2024). Associated altered biological processes include regulation of cell growth, cell senescence, cell-matrix and cell-cell adhesion, and ECM organization while key pathways include TGFβ and angiotensin signaling (Rammos et al., 2014). Because many of the characteristic features of natural aging manifest in the young Marfan aorta (Cavinato et al., 2021; Chen et al., 2023), one might expect aging to exacerbate the MFS phenotype (Gharree et al., 2022). If not, a key question becomes: What offsets key drivers of disease progression in stable Marfan aortas?

Toward this end, it is prudent first to recall characteristics of a severe ascending aortic phenotype in MFS. Although it remains unclear which transcriptional changes drive ascending aortic dissection and rupture in the *Fbn1^mgR/mgR^* MFS model, with mean survival ∼9 weeks of age (depending on breeding strategy), key changes include increases in (alphabetically) *Acan, Col12a1, Ctss, Fn1, Igfbp2, Lox, Mmp12, Serpine1, Spp1, Thbs1, and Timp1* as well as increases in *Ccl2*, *Ccr2*, *Ccl5*, *Cxcl13*, and *Itgam* but decreases in SMC contractile genes such as *Acta2* and *Myh11* (Zilberberg et al., 2015; Cikach et al., 2018; Bhushan et al., 2019; Zhang et al., 2022; Chen et al., 2023). Associated altered biological processes include chemokine signaling, ECM organization, focal adhesion, and regulation of actin cytoskeleton. Hence, dysfunctional mechanosensing and mechanoregulation of matrix likely plays a key role in early disease (Humphrey et al., 2015; Li et al., 2023) while increased inflammation may contribute to late lesion progression leading to dissection and rupture (Zhang et al., 2022; Chen et al., 2023; Kimura et al., 2024).

Prior histo-mechanical studies of the less severe *Fbn1^C1041G/+^* MFS model at 3, 6, 9, and 12 months revealed a progressive dilatation (3 to 12 mo, with possible stabilization at ∼1.33-fold) with early increases in MMP-2,9 activity (3 to 6 mo, then stable from 6 to 9 mo), progressive increases in elastin fragmentation (3 to 12 mo) and collagen deposition (6 to 12 mo), and an early decline in SM-MHC (3 to 6 mo, then stable) and late decline in SMα-A (at 12 mo), with progressive decreases in contractility (3 to 9 mo) and indications of higher circumferential stiffness and lower wall strength (Chung et al., 2007; Gharree et al., 2022). Our findings are generally consistent with these prior findings though, for the first time, we further quantified key biaxial biomechanical metrics and multi-omics while extending the age range to 2 years, which was deemed important since many manifestations of aortic aging in mice occur towards 2 years. Despite progressive increases in elastin porosity and GAGs from 12 weeks to 1 year to 2 years of age, the early increase in adventitial collagen and decrease in SMC density tended to remain similar from 1 to 2 years. So, too, the altered wall thickness, circumferential and axial material stiffness, circumferential and axial wall stress, in vivo value of axial stretch, elastic energy storage, and overall distensibility tended to remain similar from 1 to 2 years of age. Indeed, the early increase in luminal radius and decrease in contractility also tended to remain similar from 1 to 2 years. Taken together, these data suggest a stable lesion between 1 and 2 years in the *Fbn1^C1041G/+^* aorta, thus focusing attention on possible compensatory changes in the transcriptional profile and protein abundance.

Prior transcriptomic studies of the thoracic aorta from young *Fbn1^C1041G/+^* mice having progressive disease reveal many DEGs relative to age-matched WT mice. From bulk RNA-seq, these include (ordered by log_2_ fold changes for males 7 to 12 weeks of age): increased *Igfbp2*, *Serpine1*, *Tnc*, *Timp1*, *Acan*, *Tnfrsf12a*, *Itga5*, *Fn1*, *Mmp3*, *Ccn2*, *Mgp*, *Ctss* and decreased *Mylk* and *Adamts6* (Chen et al., 2021). Similarly, findings from scRNA-seq (at 4 and 24 weeks of age), included (alphabetically) increased *Bgn, Col1a1*, *Col3a1*, *Ccn2, Comp, Dcn, Loxl1, Mmp2, Serpine1, Tgfb1,* and *Thbs1* as well as a modulated SMC phenotype characterized by increased *Fn1*, *Mgp*, *Nupr1*, and *Eln* and decreased *Acta2*, *Myl9*, and *Myh11*; associated biological processes include cell proliferation and adhesion as well as collagen fibril organization (Pedroza et al., 2020). Pseudo-time trajectory analyses further suggested important temporal increases in *Ccn2*, *Igfbp2*, *Klf4, Serpine1*, *Tnfrsf11b*, and *Vcam1* and decreases in *Cnn1* and *Myh11*. In both of these studies, however, the mice were relatively young (< 6 mo). Our findings were generally consistent for young MFS mice (12 weeks), though we focused primarily on comparing results between 12 weeks and 1 year of age (during which the lesions developed) and between 1 and 2 years of age (during which the lesions appeared to be stable). We similarly found many DEGs (**Figures 4, 5, S5, S6, S8, S9; Tables S5 and S6**). Importantly, while many of these DEGs are also seen in the more severe *Fbn1^mgR/mgR^* mice (e.g., *Acan, Ccl2, Ccr2, Ctss, Fn1, Igfbp2, Itgam, Myh11, Mmp12, Serpine1, Thbs1, Tnfrs11b*), there were also distinct differences. For example, increased *Col12a1* and *Lox* were found in *Fbn1^mgR/mgR^* but not in *Fbn1^C1041G/+^* whereas decreased *Col5a1, Ltbp3,* and *Rictor* were among those found in *Fbn1^C1041G/+^*but not in *Fbn1^mgR/mgR^*. This suggests opportunities to delineate potential roles of DEGs in either promoting or preventing disease progression. Nevertheless, it is difficult to compare fold changes or sensitivities across studies.

Using consistent data collection and analysis across ages, we found statistically significant DEGs that were unique for the comparison of 12 weeks to 1 year of age but not 1 year to 2 years of age (**Table S6**), including increased *Ccr2, Lum, Mmp13,* and *Spp1* and decreased *Col15a1* and *Eln* during the first year of dilatation. Increased *Ccr2, Mmp13,* and *Spp1* are similarly found in the more severe *Fbn1^mgR/mgR^* aorta (Zhan et al., 2022; Chen et al., 2023), suggesting possible pathological roles for these transcriptional changes in particular. Conversely, statistically significant DEGs that were unique for the comparison of 1 to 2 years of age included increased *Lgals3* as well as decreased *Col5a1* and *Rictor* and a trend towards decreased *Ltbp3* at 2 years, none of which tend to be similarly differentially expressed in *Fbn1^mgR/mgR^*. Whether these altered transcripts are protective or irrelevant is unclear. Nevertheless, whereas DEGs noted here for the severe MFS phenotype associate with an inflammatory phenotype, consistent with recent reports (Chen et al., 2023; Kimura et al., 2024), those for the mild / long-term stable phenotype appear to associate with ECM development, including turnover thereof.

Finally, a prior proteomic study of the thoracic aorta from *Fbn1^C1041G/+^* mice at 10 weeks and 1 year of age similarly emphasized the importance of altered mechanosensing, particularly early alterations in fibronectin-integrin α5β1 and later alterations in vitronectin-integrin αvβ3 interactions leading to altered RICTOR and ILK signaling in the 1-year-old mice (Parker et al., 2018). Hypothesizing a role of altered TGFβ signaling, they noted increased CTGF and PAI-1 in young and older MFS aortas relative to age-matched WT controls. Altered biological pathways included integrin signaling, actin cytoskeletal remodeling, and ECM composition. Additional DPA included increased BGN, COL4A1, EMILIN1, FN, IGFBP7, MGP, THBS1, and TNC as well as multiple integrin subtypes. Our study confirmed some of these early changes while extending the age to 2 years (**Table S5**). We found altered abundance of additional proteins from 1 to 2 years, as, for example, in BGN, COL V (decreased), COL XII, COL XV, COL XVIII, FN, ILK, integrin subunits α8 and αm, LGALS3, LOXL1, LTBP3 (decreased), LUM, MGP, and THBS1, among others, though there was no detection of RICTOR. Given the long-term stability of the *Fbn1^C1041G/+^* aorta during this period, these proteins are unlikely candidates for driving aortopathy. Indeed, many of these proteins would be expected to contribute to ECM integrity, noting that our proteomics failed to detect at 2 years many proteins associated with inflammation (e.g., MMP12, 13 as well as CCL2, CCR2, CCL5, CX3CR1, CXCL13), supporting the absence of chronic inflammation in the stable *Fbn1^C1041G/+^* aortic wall.

This study is not without limitations. Unfortunately, we were unable to use some of the littermate RNA at 1 year of age, thus resulting in only two male MFS samples at that age. Given that the 12-week to 1-year period appeared to include progressive dilatation, we also compared both proteomic and transcriptomic findings for male MFS aortas (**Figure S10**) at 12 weeks + 1 year (*n* = 9 for protein and *n* = 6 for RNA, when combined) versus 2 years (*n* = 5 and 4, respectively, alone). As revealed by **Table S7**, there were statistically significant changes in both the proteomic and transcriptomic findings for only 17 of the 60 selected proteins/genes considered (28%). Of these, there was agreement (up vs. down) in 8 of the 60 pairs (13%): denoted by the gene, decreases in *Col5a1* and *Ltbp3* and increases in *Igfbp2*, *Itgam*, *Lgals3*, *Mgp*, *Thbs1*, and *Tnfrs11b* at 2 years relative to 12 week + 1 year. Among other transcripts, increased *Igfbp2*, *Itgam*, and *Thbs1* are commonly reported in *Fbn1^mgR/mgR^* aortas (Zilberberg et al., 2015; Chen et al., 2023) and *Igfbp2*, *Mgp*, *Thbs1*, and *Tnfrs11b* in young *Fbn1^C1041G/+^*aortas (Pedroza et al., 2020; Chen et al., 2021). We note further that *Adamts17*, *Ilk*, and *Rictor* were each statistically downregulated transcriptionally while the associated proteins were at such low levels that they were not detectable, hence suggesting further consistency across the two methods of assessment.

Other limitations related to the scope of the study. Vascular aging is a strong risk factor for many cardiovascular diseases in humans, including aortopathy, but the long half-life of vascular elastin (Davis, 1993) precludes full studies of vascular aging in mice (cf. Spronck et al., 2020). Indeed, even highly accelerated aging in a mouse model of progeria fails to compromise elastic laminae integrity and does not yield aortic aneurysms or dissections (Murtada et al., 2020). We focused on bulk RNA-seq consistent with our assessments of bulk rather than layer-specific mechanical properties. Future studies using single cell RNA-seq promise to provide increased insight. Similarly, new technologies enabling spatial transcriptomics and spatial biomechanical assessments (Bersi et al., 2016) promise to increase insight further. We provided histo-mechanical and transcriptional data for female and male MFS mice, but we did not seek to identify mechanisms underlying sex differences in disease severity. More attention should be directed to the observed sexual dimorphism (Cheung et al., 2017; Boczar et al., 2019). Finally, notwithstanding the availability of computational models of aortic remodeling (Humphrey, 2021), including multiscale models (Irons et al., 2021), we did not attempt to build a data-informed computational model to interpret the data further.

In summary, it is becoming increasingly apparent that DEGs secondary to *FBN1* / *Fbn1* variants can represent either pathologic consequences or protective compensations, thus suggesting a need to seek new approaches to promote compensatory mechanisms not just to prevent pathologic mechanisms. That is, there is a need to “enhance resiliency” (Goff et al., 2019). Toward this end, there is a need to understand better those mechanisms that result in stable growth of aortic aneurysms, not just disease progression. Ours is the first study to quantify together the biaxial biomechanical phenotype, multi-modality histological characterization, proteomic signature, and transcriptional profile of the Marfan aorta during periods of aortic dilatation as well as growth arrest. The histo-mechanical data suggest that the ascending aorta in the *Fbn1^C1041G/+^* mouse model of MFS worsens from 12 weeks of age to 1 year of age but remains largely stable thereafter to 2 years of age. It remains unclear whether the MFS aorta stabilizes due to temporal changes in DPA or DEGs secondary to the pathogenic variant or if processes characteristic of natural aging counter-act potentially deleterious changes due to this pathogenic variant, or both. Yet, absence of chronic inflammatory markers appears to contribute to the stability of these lesions (**Figure 4 and S5**), noting that *Ccr2, Mmp13,* and *Spp1* are highly unregulated in the more severe *Fbn1^mgR/mgR^* aortic wall (for which proteomic information is surprisingly absent) but with non-detectable associated protein abundance in the older (stable) *Fbn1^C1041G/+^* aortic wall. Compensatory changes may have included decreased LTBP3, ILK, and RICTOR and possibly PRKG1 (cf. Guo et al., 2013; Parker et al., 2018; Korneva et al., 2019), among others. It will likely be necessary to understand complex coupling effects between variants in *Fbn1* and certain DEGs given that variants compromising *Flna*, *Ilk*, and *Ltbp3* appear to promote TAAs (Shen et al., 2011; Chen et al., 2018; Guo et al., 2018) and yet decreases in *Flna*, *Ilk*, and *Ltbp3* associated herein with the stable older male *Fbn1^C1041G/+^* aortas. Toward this end, a prior study showed an improved aortic phenotype in *Fbn1^mgR/mgR^ ; Ltbp3^-/-^* mice (Korneva et al., 2019), consistent with the present finding of decreased *Ltbp3* / LTBP-3 in the older *Fbn1^C1041G/+^* mice. Although much remains to be learned, it appears that targeting inflammation should be explored to prevent late-stage disease in the Marfan aorta as should methods to increase smooth muscle contractility, increase ECM cross-linking, and decrease mTOR signaling.

## ACKNOWLEDGMENTS

This work was supported by grants from the US National Institutes of Health (R01 HL168473 to R. Assi, R01 HL169147 and P01 HL169168 to JDH, and R03 AG074063 to EPM for the aging controls) and the Leducq Foundation (erADDicate). We acknowledge early technical support from Drs. Justyna Niestrawska and Maria Jesus Ruiz-Rodriguez as well as assistance with MS sample preparation and data collection by Florine Collin and Dr. Weiwei Wang. We thank the Keck MS and Proteomics Resource at Yale School of Medicine for providing the necessary mass spectrometers and biotechnology tools funded, in part, by the Yale School of Medicine and the US National Institutes of Health (S10OD02365101, S10OD019967, and S10OD018034).

## DATA AVAILABILITY

Geometric and biomechanical data are tabulated in Supplementary Materials. The mass spectrometry proteomics data have been deposited to the ProteomeXchange Consortium via the PRIDE partner repository and will be made available with the dataset identifier PXD061014. The transcriptomic data will be uploaded to NIH GEO upon acceptance.

## REFERENCES

1. Bellini C, Korneva A, Zilberberg L, Ramirez F, Rifkin DB, Humphrey JD. Differential ascending and descending aortic mechanics parallel aneurysmal propensity in a mouse model of Marfan syndrome. J Biomech. 2016;49(12):2383–2389.

2. Bersi MR, Bellini C, Di Achille P, Humphrey JD, Genovese K, Avril S. Novel methodology for characterizing regional variations in the material properties of murine aortas. J Biomech Eng. 2016;138:0710051–07100515.

3. Bhushan R, Altinbas L, Jäger M, Zaradzki M, Lehmann D, Timmermann B, Clayton NP, Zhu Y, Kallenbach K, Kararigas G, Robinson PN. An integrative systems approach identifies novel candidates in Marfan syndrome-related pathophysiology. J Cell Mol Med. 2019;23(4):2526–2535.

4. Boczar KE, Cheung K, Boodhwani M, Beauchesne L, Dennie C, Nagpal S, Chan K, Coutinho T. Sex differences in thoracic aortic aneurysm growth. Hypertension. 2019;73:190–196.

5. Bordner KA, George ED, Carlyle BC, Duque A, Kitchen RR, Lam TT, Colangelo CM, Stone KL, Abbott TB, Mane SM, Nairn AC, Simen AA. Functional genomic and proteomic analysis reveals disruption of myelin-related genes and translation in a mouse model of early life neglect. Front Psychiatry 2011; 2:18.

6. Busnadiego O, Gorbenko Del Blanco D, González-Santamaría J, Habashi JP, Calderon JF, Sandoval P, Bedja D, Guinea-Viniegra J, Lopez-Cabrera M, Rosell-Garcia T, Snabel JM, Hanemaaijer R, Forteza A, Dietz HC, Egea G, Rodriguez-Pascual F. Elevated expression levels of lysyl oxidases protect against aortic aneurysm progression in Marfan syndrome. J Mol Cell Cardiol. 2015;85:48–57.

7. Cavinato C, Chen M, Weiss D, Rodriguez-Ruiz MJ, Schwartz MA, Humphrey JD. Progressive microstructural deterioration dictates evolving biomechanical dysfunction in the Marfan aorta. Front Cardiovasc Med 2021;8:800730.

8. Chen MH, Choudhury S, Hirata M, Khalsa S, Chang B, Walsh CA. Thoracic aortic aneurysm in patients with loss of function Filamin A mutations: Clinical characterization, genetics, and recommendations. Am J Med Genet A. 2018;176(2):337–350.

9. Chen JZ, Sawada H, Ye D, Katsumata Y, Kukida M, Ohno-Urabe S, Moorleghen JJ, Franklin MK, Howatt DA, Sheppard MB, Mullick AE, Lu HS, Daugherty A. Deletion of AT1a (angiotensin ii type 1a) receptor or inhibition of angiotensinogen synthesis attenuates thoracic aortopathies in Fibrillin1(C1041G/+) mice. Arterioscler Thromb Vasc Biol. 2021;41(10):2538–2550.

10. Chen M, Cavinato C, Hansen J, Tanaka K, Ren P, Hassab A, Li DS, Joshuao E, Tellides G, Iyengar R, Humphrey JD, Schwartz MA. Fibronectin-integrin alpha5 signaling promotes thoracic aortic aneurysm in a mouse model of Marfan syndrome. ATVB 2023;43:e132–150.

11. Cheung K, Boodhwani M, Chan KL, Beauchesne L, Dick A, Coutinho T. Thoracic aortic aneurysm growth: role of sex and aneurysm etiology. J Am Heart Assoc. 2017;6:e003792.

12. Chung AW, Au Yeung K, Sandor GG, Judge DP, Dietz HC, van Breemen C. Loss of elastic fiber integrity and reduction of vascular smooth muscle contraction resulting from the upregulated activities of matrix metalloproteinase-2 and -9 in the thoracic aortic aneurysm in Marfan syndrome. Circ Res. 2007;101:512–22.

13. Cikach FS, Koch CD, Mead TJ, Galatioto J, Willard BB, Emerton KB, Eagleton MJ, Blackstone EH, Ramirez F, Roselli EE, Apte SS. Massive aggrecan and versican accumulation in thoracic aortic aneurysm and dissection. JCI Insight. 2018;3(5):e97167.

14. Cook JR, Carta L, Galatioto J, Ramirez F. Cardiovascular manifestations in Marfan syndrome and related diseases; multiple genes causing similar phenotypes. Clin Genet. 2015;87:11–20.

15. Cyron C, Wilson JS, Humphrey JD (2014) Mechanobiological stability: A new paradigm to understand the enlargement of aneurysms. J R Soc Interface 11: 20140680.

16. Davis EC. Stability of elastin in the developing mouse aorta: a quantitative radioautographic study. Histochemistry. 1993;100:17–26.

17. Deleeuw V, De Clercq A, De Backer J, Sips P. An Overview of Investigational and Experimental Drug Treatment Strategies for Marfan Syndrome. J Exp Pharmacol. 2021;13:755–779.

18. de Wit A, Vis K, Jeremy RW. Aortic stiffness in heritable aortopathies: relationship to aneurysm growth rate. Heart Lung Circ. 2013;22:3–11.

19. Eberth JF, Taucer AI, Wilson E, Humphrey JD. Mechanics of carotid arteries in a mouse model of Marfan Syndrome. Ann Biomed Eng. 2009;37:1093–104.

20. Ferruzzi J, Bersi MR, Humphrey JD. Biomechanical phenotyping of central arteries in health and disease: advantages of and methods for murine models. Ann Biomed Eng. 2013;41(7):1311–30.

21. Ferruzzi J, Madziva D, Caulk AW, Tellides G, Humphrey JD. Compromised mechanical homeostasis in arterial aging and associated cardiovascular consequences. Biomech Model Mechanobiol. 2018;17(5):1281–1295.

22. Greenbaum D, Colangelo C, Williams K, Gerstein M. Comparing protein abundance and mRNA expression levels on a genomic scale. Genome Biol. 2003;4:1–8.

23. Gharraee N, Sun Y, Swisher JA, Lessner SM. Age and sex dependency of thoracic aortopathy in a mouse model of Marfan syndrome. Am J Physiol Heart Circ Physiol. 2022;322:H44–H56

24. Gomez D, Kessler K, Borges LF, Richard B, Touat Z, Ollivier V, Mansilla S, Bouton MC, Alkoder S, Nataf P, Jandrot-Perrus M, Jondeau G, Vranckx R, Michel JB. Smad2-dependent protease nexin-1 overexpression differentiates chronic aneurysms from acute dissections of human ascending aorta. Arterioscler Thromb Vasc Biol. 2013;33:2222–32.

25. Goff DC Jr, Buxton DB, Pearson GD, Wei GS, Gosselin TE, Addou EA, Stoney CM, Desvigne-Nickens P, Srinivas PR, Galis ZS, Pratt C, Kit KBK, Maric-Bilkan C, Nicastro HL, Wong RP, Sachdev V, Chen J, Fine L, Writing Group for the Division of Cardiovascular Sciences’ Strategic Vision Implementation Plan. Implementing the National Heart, Lung, and Blood Institute’s strategic vision in the Division of Cardiovascular Sciences. Circ Res. 2019;124(4):491–497.

26. Guo DC, Regalado E, Casteel DE, Santos-Cortez RL, Gong L, Kim JJ, Dyack S, Horne SG, Chang G, Jondeau G, Boileau C, Coselli JS, Li Z, Leal SM, Shendure J, Rieder MJ, Bamshad MJ, Nickerson DA; GenTAC Registry Consortium; National Heart, Lung, and Blood Institute Grand Opportunity Exome Sequencing Project; Kim C, Milewicz DM. Recurrent gain-of-function mutation in PRKG1 causes thoracic aortic aneurysms and acute aortic dissections. Am J Hum Genet. 2013; 93(2):398–404.

27. Guo DC, Regalado ES, Pinard A, Chen J, Lee K, Rigelsky C, Zilberberg L, Hostetler EM, Aldred M, Wallace SE, Prakash SK; University of Washington Center for Mendelian Genomics; Leal SM, Bamshad MJ, Nickerson DA, Natowicz M, Rifkin DB, Milewicz DM. LTBP3 pathogenic variants predispose individuals to thoracic aortic aneurysms and dissections. Am J Hum Genet. 2018;102(4):706–712.

28. Hirosawa M, Hoshida M, Ishikawa M, Toya T. MASCOT: multiple alignment system for protein sequences based on three-way dynamic programming. Comput Appl Biosci. 1993;9(2):161-7.

29. Humphrey JD, Schwartz MA, Tellides G, Milewicz DM. Role of mechanotransduction in vascular biology: focus on thoracic aortic aneurysms and dissections. Circ Res. 2015;116(8):1448–61.

30. Humphrey JD. Constrained mixture models of tissue growth and remodeling – Twenty years after. J Elasticity 2021;145:49–75.

31. Irons L, Latorre M, Humphrey JD. From transcript to tissue: multiscale modeling from cell signaling to matrix remodeling. Ann Biomed Eng. 2021;49(7):1701–1715.

32. Judge DP, Biery NJ, Keene DR, Geubtner J, Myers L, Huso DL, Sakai LY, Dietz HC. Evidence for a critical contribution of haploinsufficiency in the complex pathogenesis of Marfan syndrome. J Clin Invest. 2004;114:172–81.

33. Kawamura Y, Murtada S-I, Tellides G, Humphrey JD. Adventitial remodeling protects against aortic rupture following late smooth-muscle-specific disruption of TGFβ signaling. J Mech Behav Biomed Matl 2021;116: 104264.

34. Kelleher CM, McLean SE, Mecham RP. Vascular extracellular matrix and aortic development. Curr Top Dev Biol. 2004;62:153–88.

35. Kimura K, Motoyama E, Kanki S, Asano K, Sheikh MAA, Clarin MTRDC, Raja E, Takeda M, Ishii R, Murata K, Deleeuw V, Sips P, Mosquera LM, De Backer J, Mizuno S, Sakai LY, Nakamura T, Yanagisawa H. A novel genetic mouse model of fatal aortic dissection reveals massive inflammatory cell infiltration in the thoracic aorta. bioRxiv doi: 10.1101/2024.05.02.592287.

36. Korneva A, Zilberberg L, Rifkin DB, Humphrey JD, Bellini C. Absence of LTBP-3 attenuates the aneurysmal phenotype but not spinal effects on the aorta in Marfan syndrome. Biomech Model Mechanobiol. 2019;18:261–273.

37. Li DS, Cavinato C, Latorre M, Humphrey JD. Computational modeling distinguished diverse contributors to aneurysmal progression in the Marfan aorta. Proceed R Soc Lond A 2023; 479: 20230116.

38. Milewicz DM, Braverman AC, De Backer J, Morris SA, Boileau C, Maumenee IH, Jondeau G, Evangelista A, Pyeritz RE. Marfan syndrome. Nat Rev Dis Primers. 2021;7(1):64.

39. Nollen GJ, Groenink M, Tijssen JG, Van Der Wall EE, Mulder BJ. Aortic stiffness and diameter predict progressive aortic dilatation in patients with Marfan syndrome. Eur Heart J. 2004;25:1146–52.

40. Parker SJ, Stotland A, MacFarlane E, Wilson N, Orosco A, Venkatraman V, Madrid K, Gottlieb R, Dietz HC, Van Eyk JE. Proteomics reveals Rictor as a noncanonical TGF-β signaling target during aneurysm progression in Marfan mice. Am J Physiol Heart Circ Physiol. 2018;315:H1112–H1126.

41. Pedroza AJ, Tashima Y, Shad R, Cheng P, Wirka R, Churovich S, Nakamura K, Yokoyama N, Cui JZ, Iosef C, Hiesinger W, Quertermous T, Fischbein MP. Single-Cell transcriptomic profiling of vascular smooth muscle cell phenotype modulation in Marfan syndrome aortic aneurysm. Arterioscler Thromb Vasc Biol. 2020;40(9):2195–2211.

42. Pereira L, Lee SY, Gayraud B, Andrikopoulos K, Shapiro SD, Bunton T, Biery NJ, Dietz HC, Sakai LY, Ramirez F. Pathogenetic sequence for aneurysm revealed in mice underexpressing fibrillin-1. Proc Natl Acad Sci U S A. 1999;96:3819–23.

43. Murtada SI, Kawamura Y, Caulk AW, Ahmadzadeh H, Mikush N, Zimmerman K, Kavanagh D, Weiss D, Latorre M, Zhuang ZW, Shadel GS, Braddock DT, Humphrey JD. Paradoxical aortic stiffening and subsequent cardiac dysfunction in Hutchinson-Gilford progeria syndrome. J R Soc Interface. 2020;17(166):20200066.

44. Rammos C, Hendgen-Cotta UB, Deenen R, Pohl J, Stock P, Hinzmann C, Kelm M, Rassaf T. Age-related vascular gene expression profiling in mice. Mech Ageing Dev. 2014;135:15–23.

45. Rivera CF, Farra YM, Silvestro M, Medvedovsky S, Matz J, Pratama MY, Vlahos J, Ramkhelawon B, Bellini C. Mapping the unicellular transcriptome of the ascending thoracic aorta to changes in mechanosensing and mechanoadaptation during aging. Aging Cell. 2024;23(8):e14197.

46. Roccabianca S, Bellini C, Humphrey JD. Computational modelling suggests good, bad and ugly roles of glycosaminoglycans in arterial wall mechanics and mechanobiology. J R Soc Interface. 2014;11(97):20140397.

47. Selamet Tierney ES, Levine JC, Sleeper LA, Roman MJ, Bradley TJ, Colan SD, Chen S, Campbell MJ, Cohen MS, De Backer J, Heydarian H, Hoskoppal A, Lai WW, Liou A, Marcus E, Nutting A, Olson AK, Parra DA, Pearson GD, Pierpont ME, Printz BF, Pyeritz RE, Ravekes W, Sharkey AM, Srivastava S, Young L, Lacro RV; Pediatric Heart Network Investigators. Influence of aortic stiffness on aortic-root growth rate and outcome in patients with the Marfan Syndrome. Am J Cardiol. 2018;121:1094–1101.

48. Schwill S, Seppelt P, Grünhagen J, Ott CE, Jugold M, Ruhparwar A, Robinson PN, Karck M, Kallenbach K. The fibrillin-1 hypomorphic mgR/mgR murine model of Marfan syndrome shows severe elastolysis in all segments of the aorta. J Vasc Surg. 2013;57(6):1628–36, 1636.e1-3.

49. Shen D, Li J, Lepore JJ, Anderson TJ, Sinha S, Lin AY, Cheng L, Cohen ED, Roberts JD Jr, Dedhar S, Parmacek MS, Gerszten RE. Aortic aneurysm generation in mice with targeted deletion of integrin-linked kinase in vascular smooth muscle cells. Circ Res. 2011;109(6):616–28.

50. Spronck B, Ferruzzi J, Bellini C, Caulk AW, Murtada SI, Humphrey JD. Aortic remodeling is modest and sex-independent in mice when hypertension is superimposed on aging. J Hypertens. 2020;38(7):1312–1321.

51. Sun Y, Asano K, Sedes L, Cantalupo A, Hansen J, Iyengar R, Walsh MJ, Ramirez F. Dissecting aortic aneurysm in Marfan syndrome is associated with losartan-sensitive transcriptomic modulation of aortic cells. JCI Insight. 2023;8:e168793.

52. Weiss D, Rego BV, Cavinato C, Li DS, Kawamura Y, Emuna N, Humphrey JD. Effects of age, sex, and extracellular matrix integrity on aortic dilatation and rupture in a mouse model of Marfan Syndrome. Arterioscler Thromb Vasc Biol. 2023; 43: e358–372.

53. Weiss D, Yeung N, Ramachandra AB, Humphrey JD. Transcriptional regulation of postnatal aortic development. Cells Dev. 2024;180:203971.

54. Westenberg JJ, Scholte AJ, Vaskova Z, van der Geest RJ, Groenink M, Labadie G, van den Boogaard PJ, Radonic T, Hilhorst-Hofstee Y, Mulder BJ, Kroft LJ, Reiber JH, de Roos A. Age-related and regional changes of aortic stiffness in the Marfan syndrome: assessment with velocity-encoded MRI. J Magn Reson Imaging. 2011;34:526–31.

55. Wilson JS, Baek S, Humphrey JD. Parametric study of effects of collagen turnover on the natural history of abdominal aortic aneurysms. Proc Math Phys Eng Sci. 2013;469(2150):20120556.

56. Wu J, Montaniel KR, Saleh MA, Xiao L, Chen W, Owens GK, Humphrey JD, Majesky MW, Paik DT, Hatzopoulos AK, Madhur MS, Harrison DG. Origin of matrix-producing cells that contribute to aortic fibrosis in hypertension. Hypertension. 2016;67(2):461–8

57. Yamashiro Y, Thang BQ, Shin SJ, Lino CA, Nakamura T, Kim J, Sugiyama K, Tokunaga C, Sakamoto H, Osaka M, Davis EC, Wagenseil JE, Hiramatsu Y, Yanagisawa H. Role of Thrombospondin-1 in mechanotransduction and development of thoracic aortic aneurysm in mouse and humans. Circ Res. 2018;123(6):660–672.

58. Zhang RM, Tiedemann K, Muthu ML, Dinesh NEH, Komarova S, Ramkhelawon B, Reinhardt DP. Fibrillin-1-regulated miR-122 has a critical role in thoracic aortic aneurysm formation. Cell Mol Life Sci. 2022;79(6):314.

59. Zilberberg L, Phoon CK, Robertson I, Dabovic B, Ramirez F, Rifkin DB. Genetic analysis of the contribution of LTBP-3 to thoracic aneurysm in Marfan syndrome. Proc Natl Acad Sci U S A. 2015;112:14012–7.

