## Supplemental material for "Growth Arrest of Thoracic Aortic Aneurysms in Aging Marfan Mice"

**Tables and Figures**

**Table S1**. Best-fit values of the material parameters for passive biaxial mechanical behaviors of the ascending thoracic aorta (ATA) from all 12 groups: female (F) and male (M), wild-type (WT) and Marfan syndrome (MFS) at 12 weeks, 1 year, and 2 years of age.

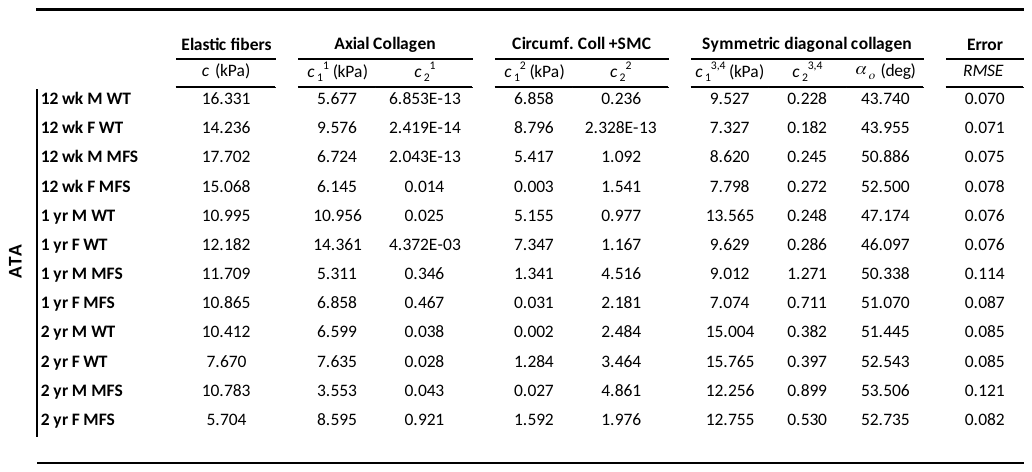

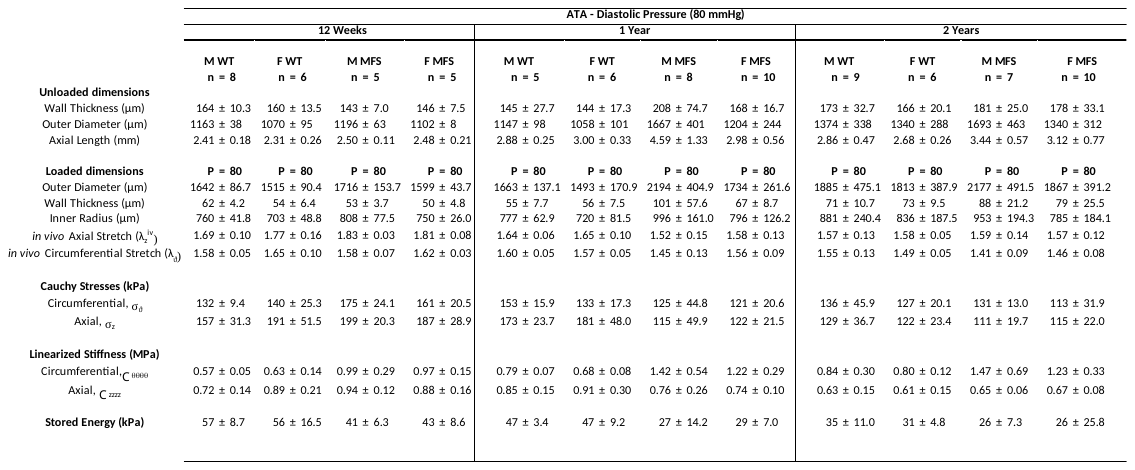
**Table S2**. Measured and calculated biomechanical metrics for all 12 groups of mice, with load-dependent values computed at a common diastolic pressure (80 mmHg) but individual in vivo values of axial stretch.

**Table S3**. Measured and calculated biomechanical metrics for all 12 groups of mice, with load-dependent values computed at a common pressure of 100 mmHg but individual in vivo values of axial stretch.

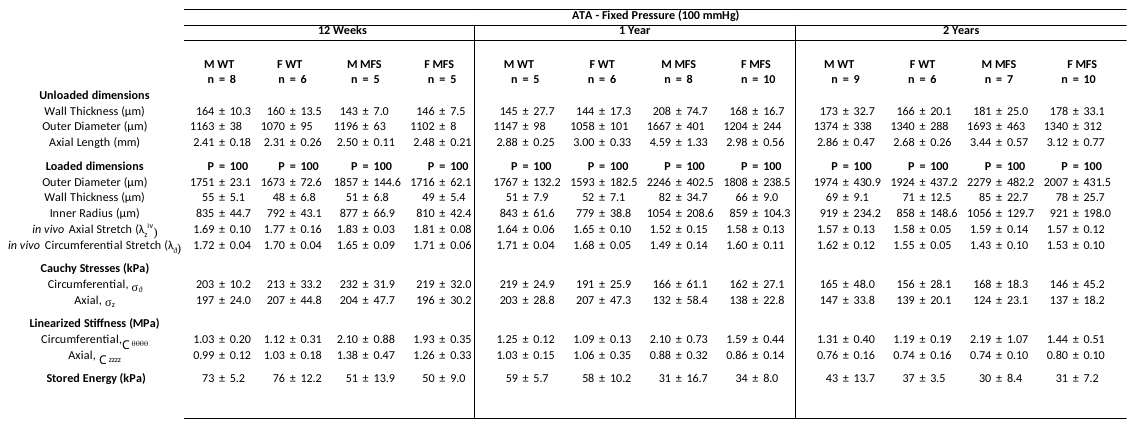

**Table S4**. Measured and calculated biomechanical metrics for all 12 groups of mice, with load-dependent values computed at a common systolic pressure (120 mmHg) but individual in vivo values of axial stretch.

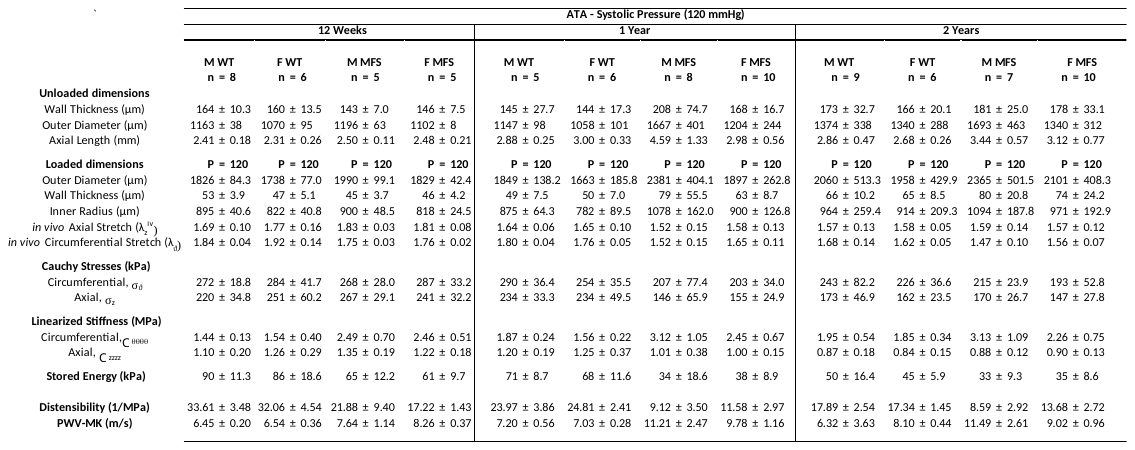

**Table S5**. Statistical comparisons based on proteomics for 60 select proteins for male MFS mice across three comparisons: 12 weeks vs. 1 year of age, 1 vs. 2 years of age, and 12 weeks vs. 2 years of age. Pink cells highlight statistical significance (*p* < 0.05).

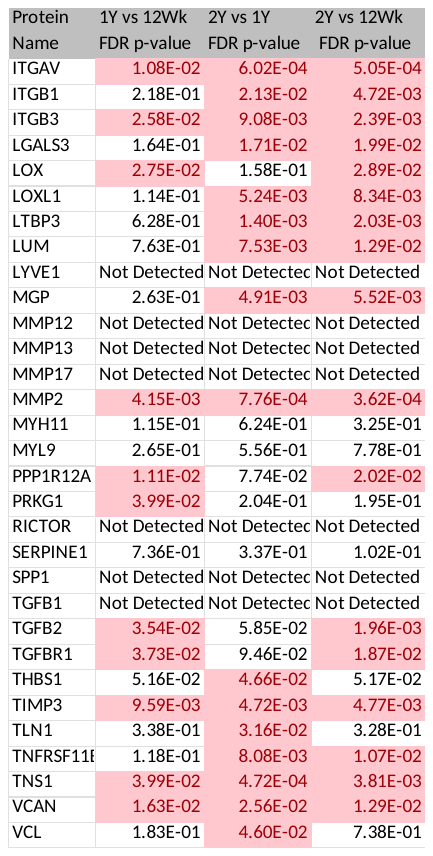

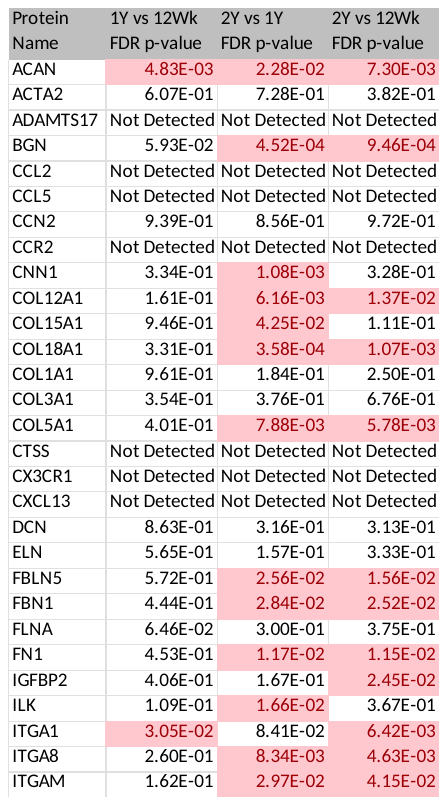

**Table S6**. Statistical comparisons based on transcriptomics for 60 select genes for male MFS mice across three comparisons: 12 weeks vs. 1 year of age, 1 vs. 2 years of age, and 12 weeks vs. 2 years of age. Pink cells highlight statistical significance (*p* < 0.05).

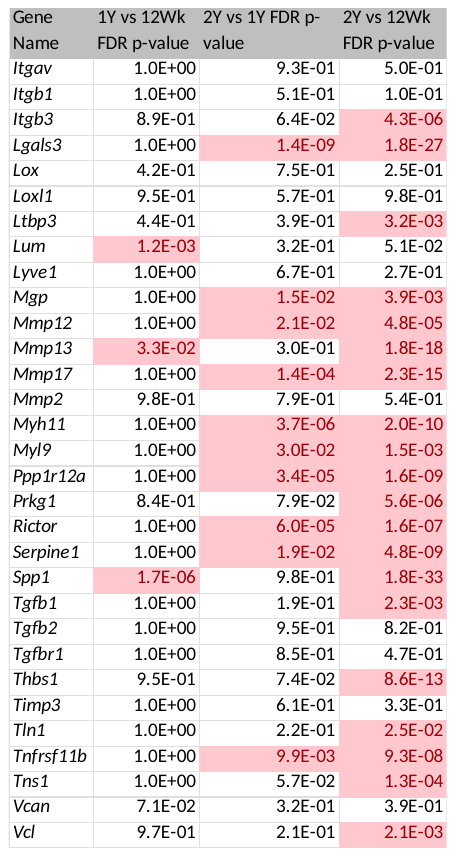

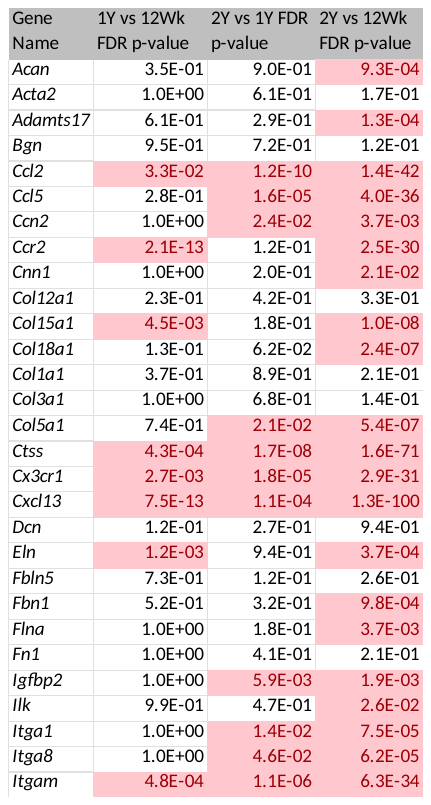

**Table S7**. Comparisons for 60 select proteins (left) and 60 select genes (right) for male MFS mice based on data for 12 weeks + 1 year of age versus those at 2 years of age alone.

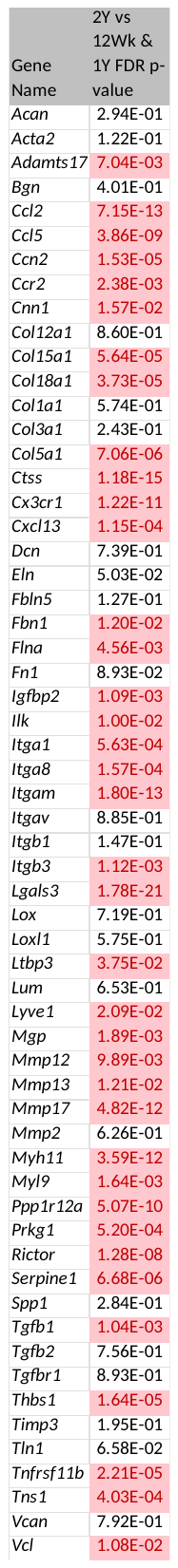

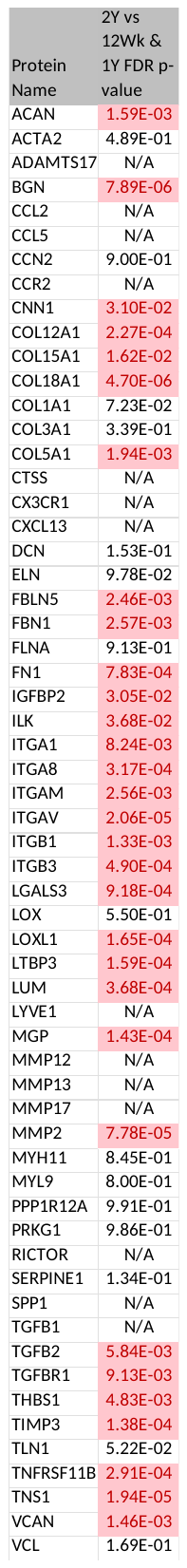

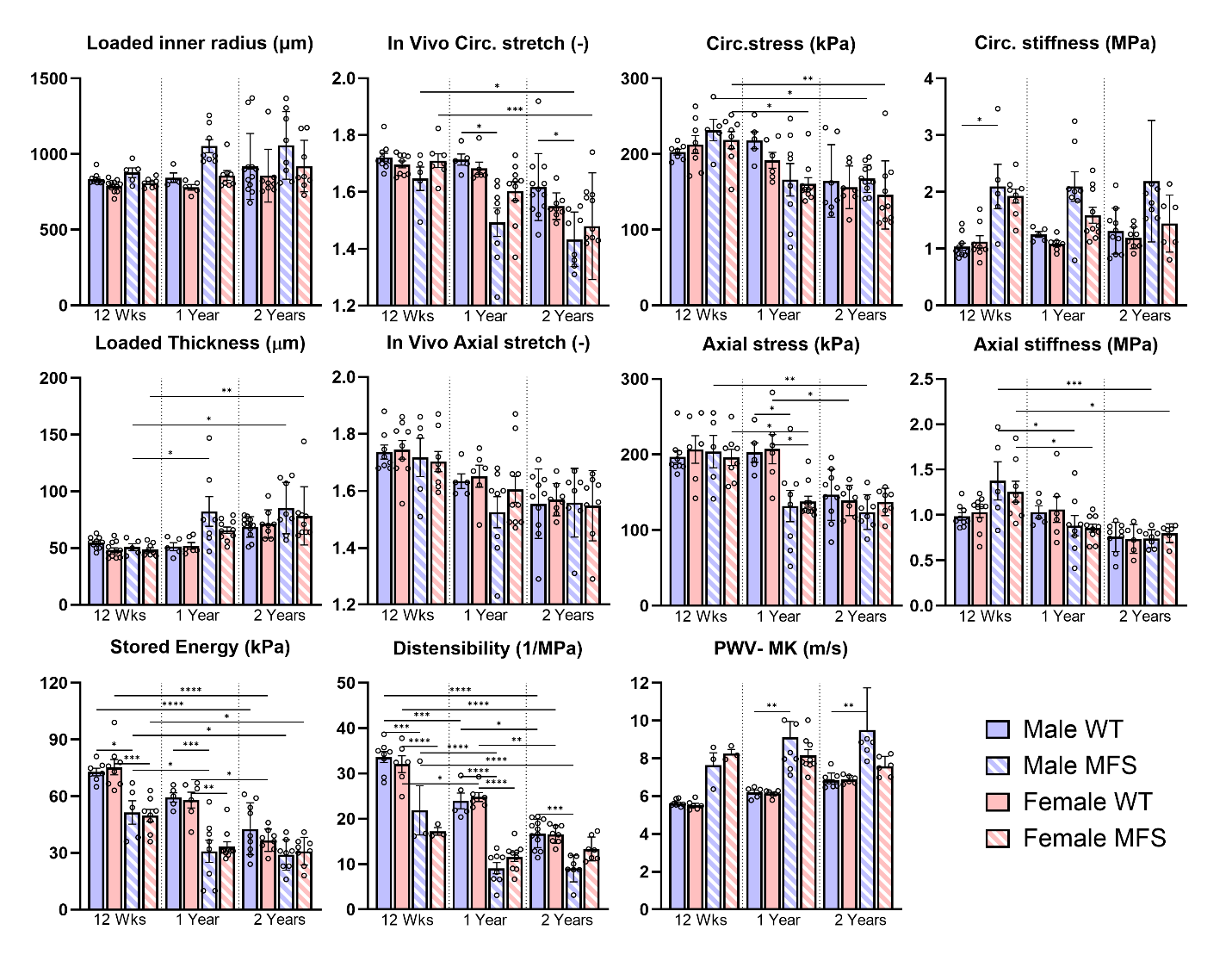

**Figure S1**. Biomechanical metrics for both sexes (female – red, male - blue), both genotypes (WT – solid, MFS – cross-hatched), and all three ages (12 weeks, 1 year, 2 years) shown at a common pressure of 100 mmHg but specimen-specific in vivo stretch. Values were calculated using material parameters in Supplemental Table S1. Mean ± SEM, with *, **, ***, **** denoting significance at *p* < 0.05, 0.01, 0.001, 0.0001, respectively. See Tables S2-S4 for tabulated values at three different pressures.

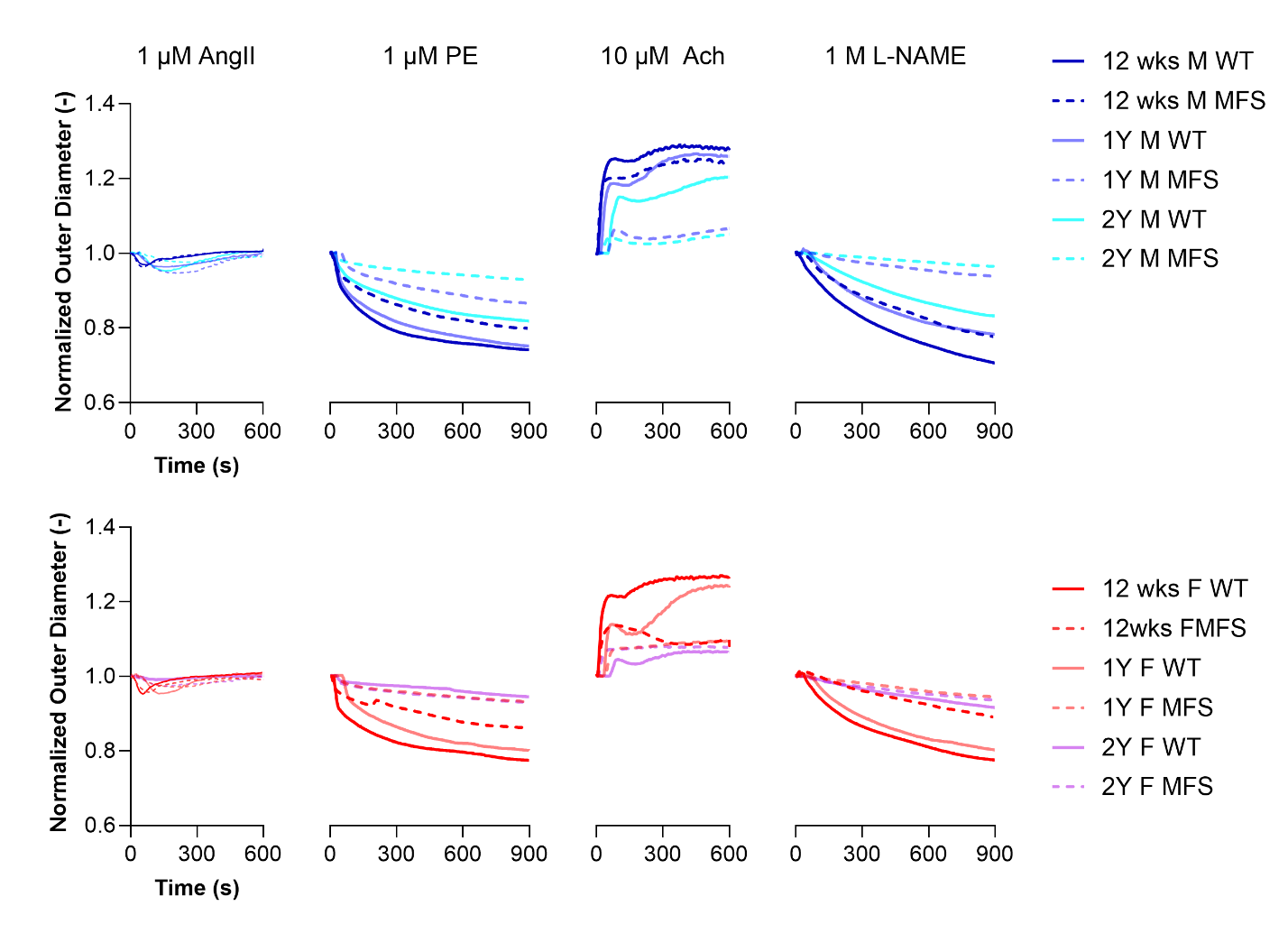

**Figure S2**. Illustrative reductions in outer dimeter for male (blue - top) and female (red - bottom), WT (solid) and MFS (dot-dashed) aortas subjected to four different vasoactive substances: angiotensin II (AngII), phenylephrine (PE), and the endothelial-derived nitric oxide synthase-inhibitor L-NAME at indicated dosages for 15 minutes, and acetylcholine (Ach) at the indicated dose for 10 minutes.

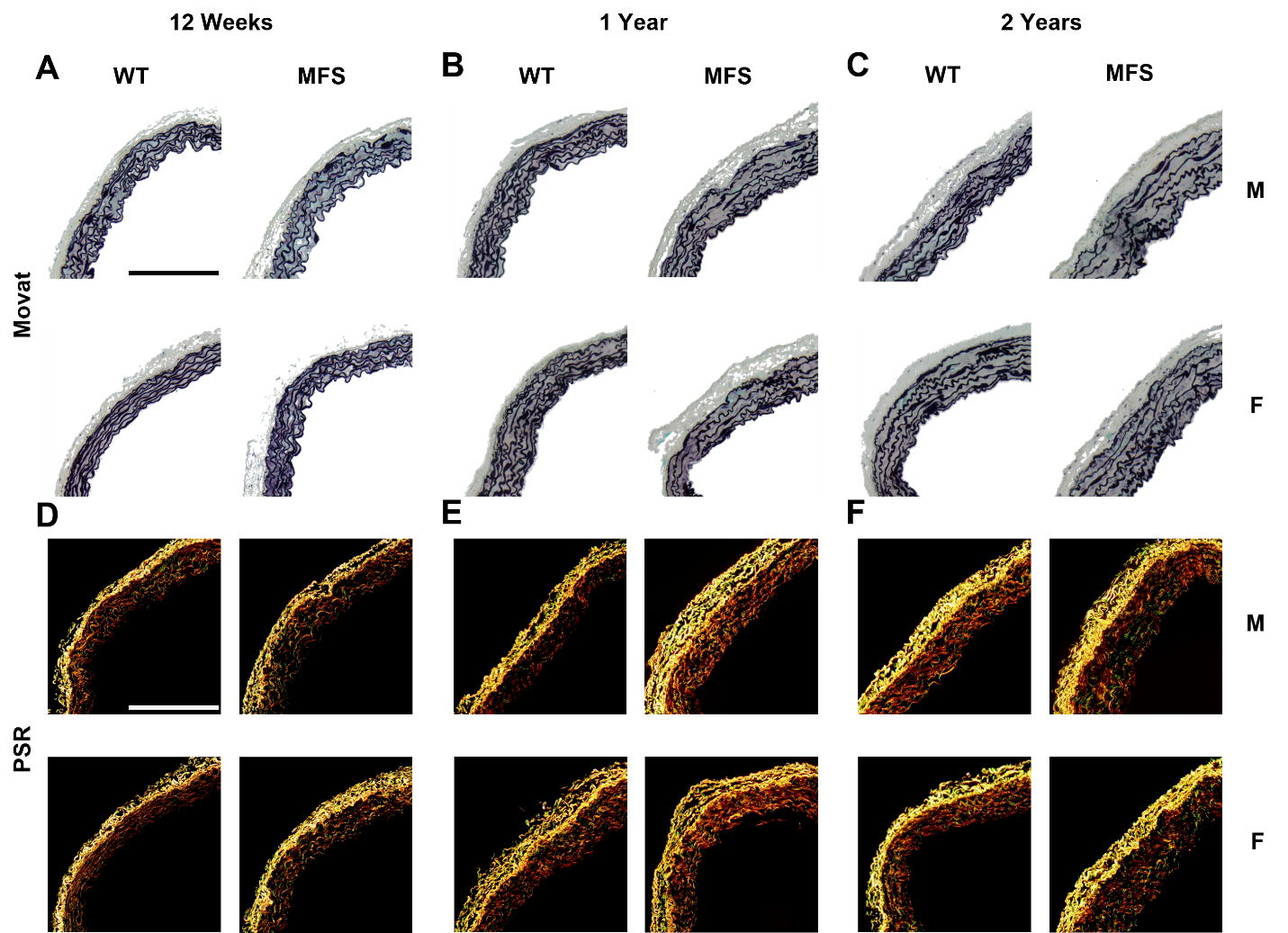

**Figure S3**. Similar to Figure 2 in the main text, but additional representative histological images for male (M) and female (F), wild-type (WT) and Marfan syndrome (MFS) aortas at all three ages: 12 weeks, 1 year, and 2 years. A-C. Bright field light microscopic images of Movat-stained sections: elastin (black), collagen (grey/yellow), glycosaminoglycans (blue), and cytoplasm (pink/red). D-F. Polarized light microscopic images of collagen from picrosirius red (PSR) stained sections. Scale bar = 100 µm.

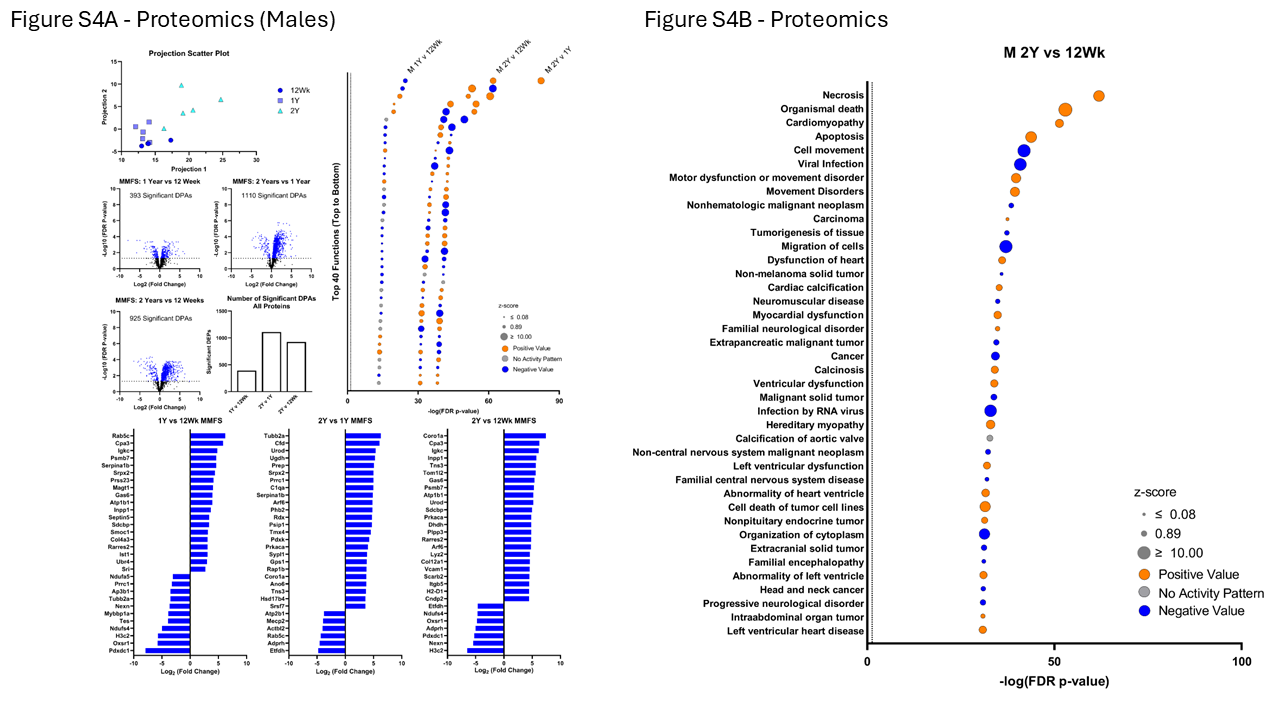

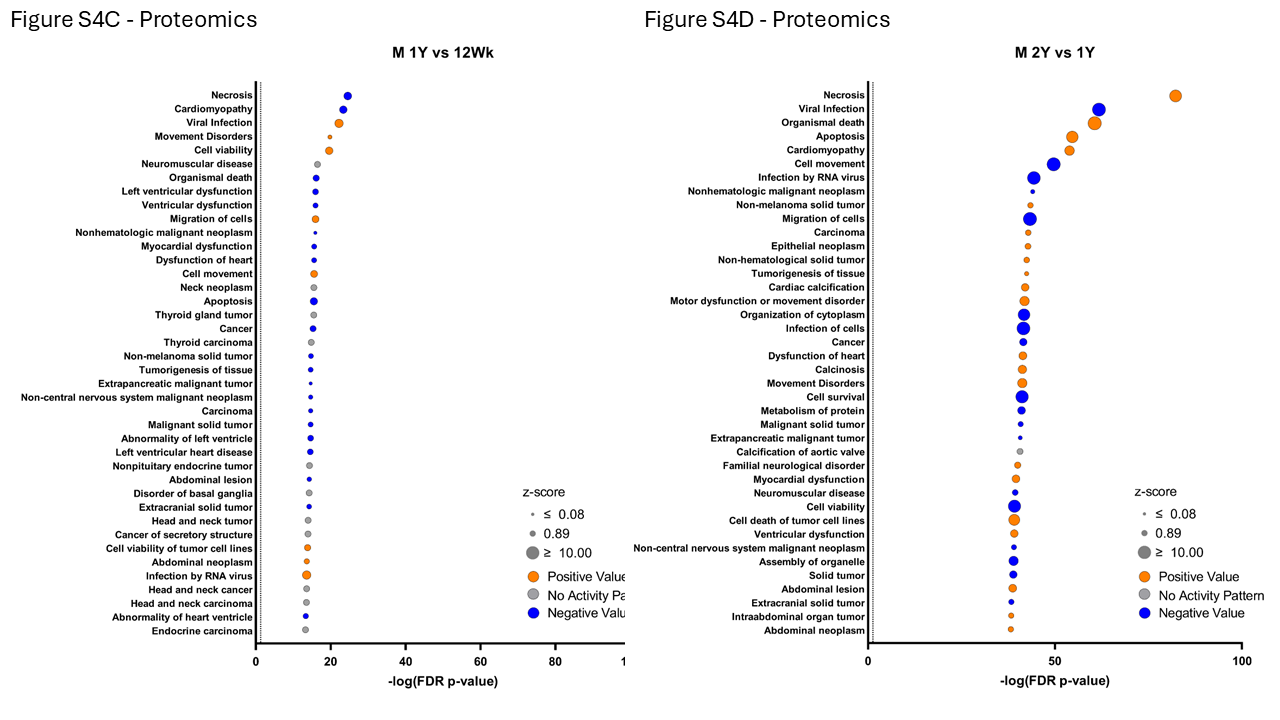

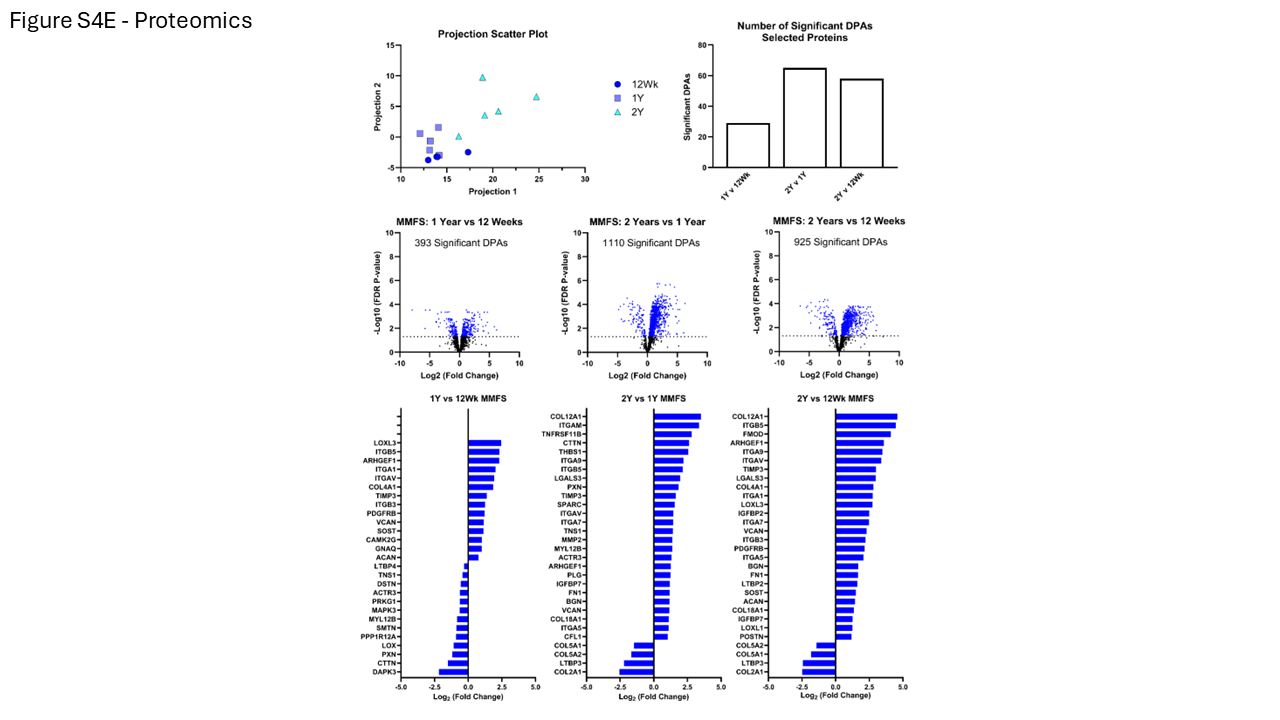

**Figure S4**. Differential protein abundance (DPA) across all three ages of male MFS aortas. A. Principal component analysis, volcano plots, and log2 fold changes. B. Associated biological processes for 2 years vs. 12 weeks of age. C. Associated biological processes for 1 year vs. 12 weeks of age. D. Associated biological processes for 2 years vs. 1 year of age. E. Similar to panel A except for select proteins.

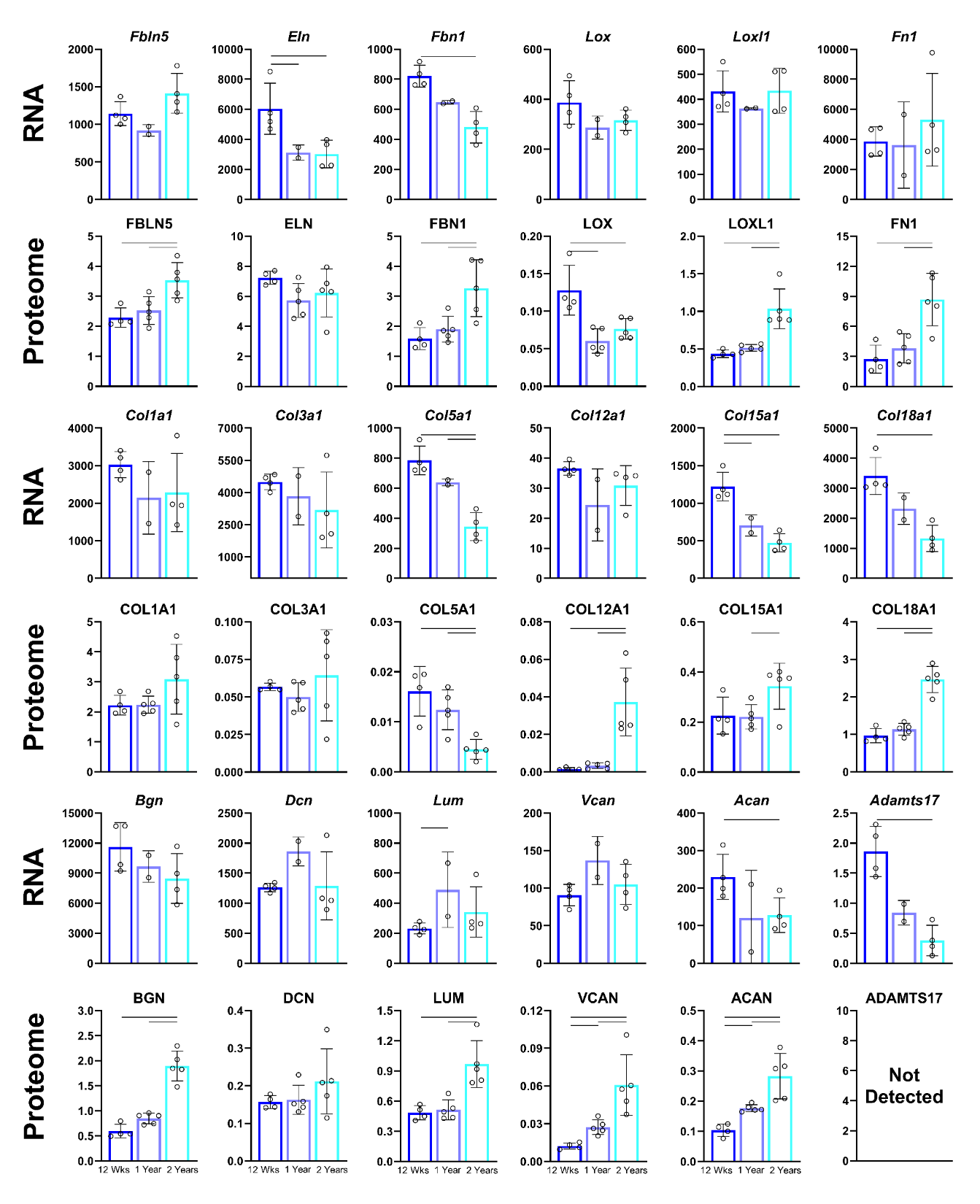

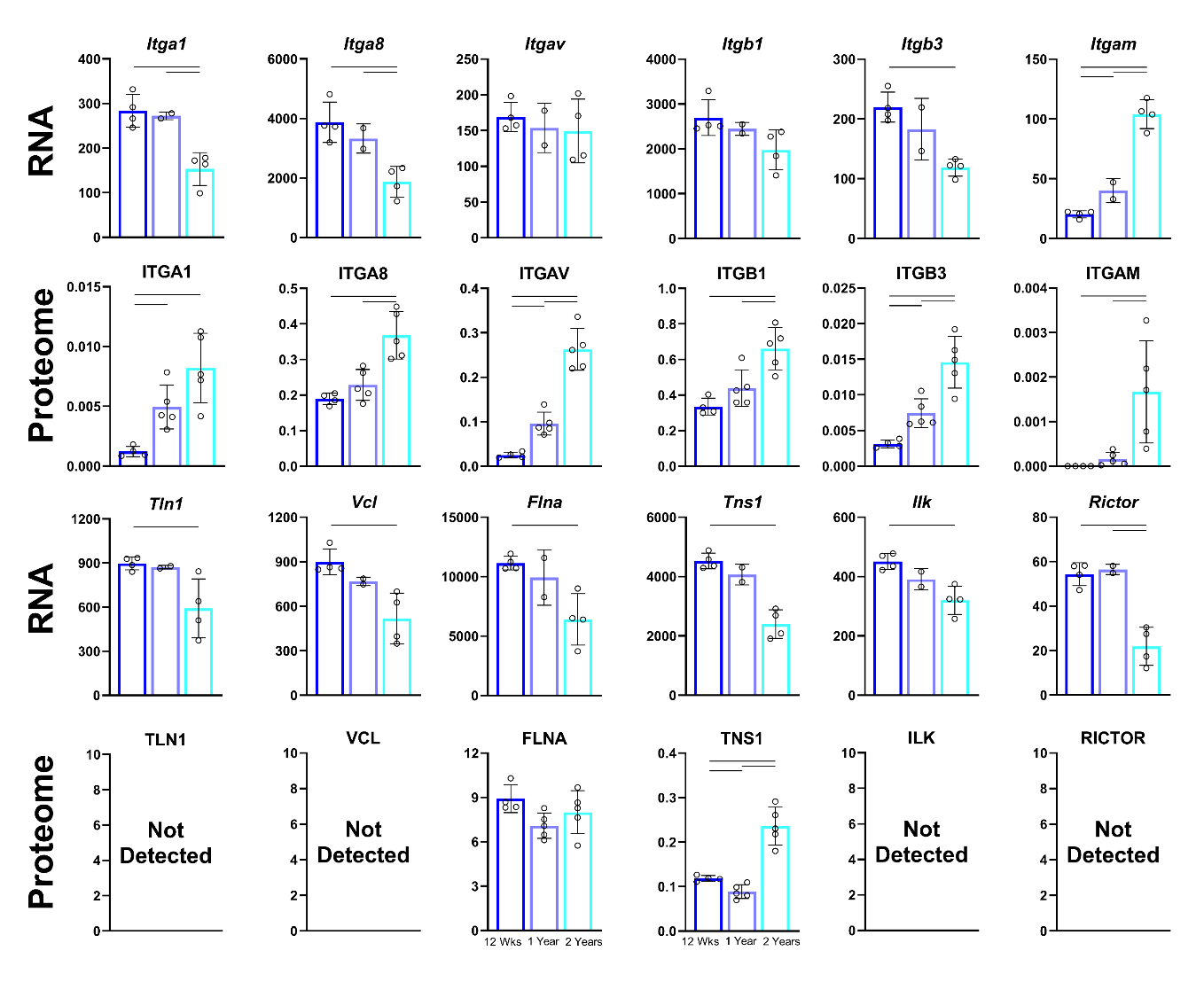

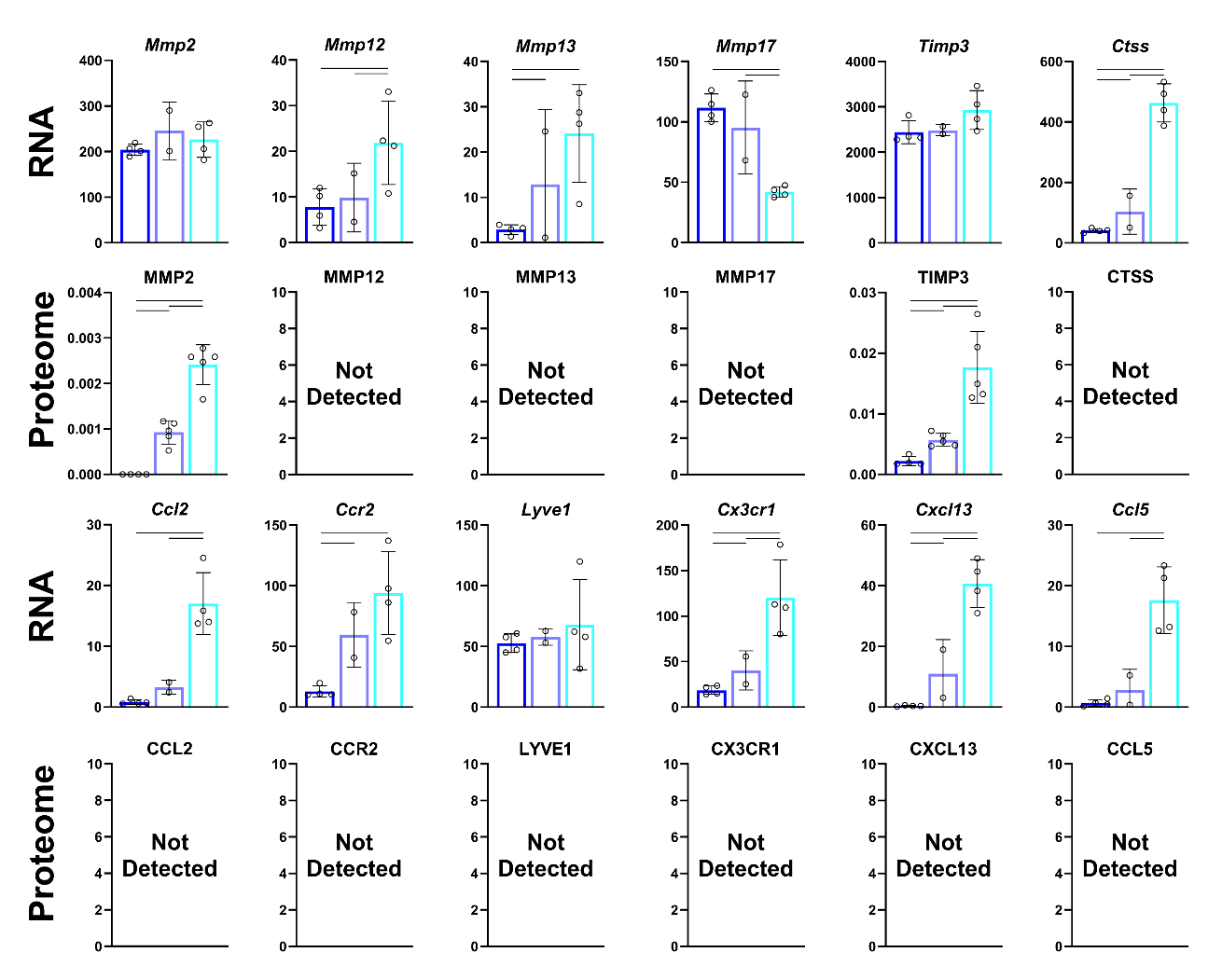

**Figure S5**. Similar for Figure 4 but for additional transcripts (RNA) and proteins (proteome), collectively totally 60. In each panel, left bar (12 weeks of age), middle bar (1 year of age), and right bar (2 years of age).

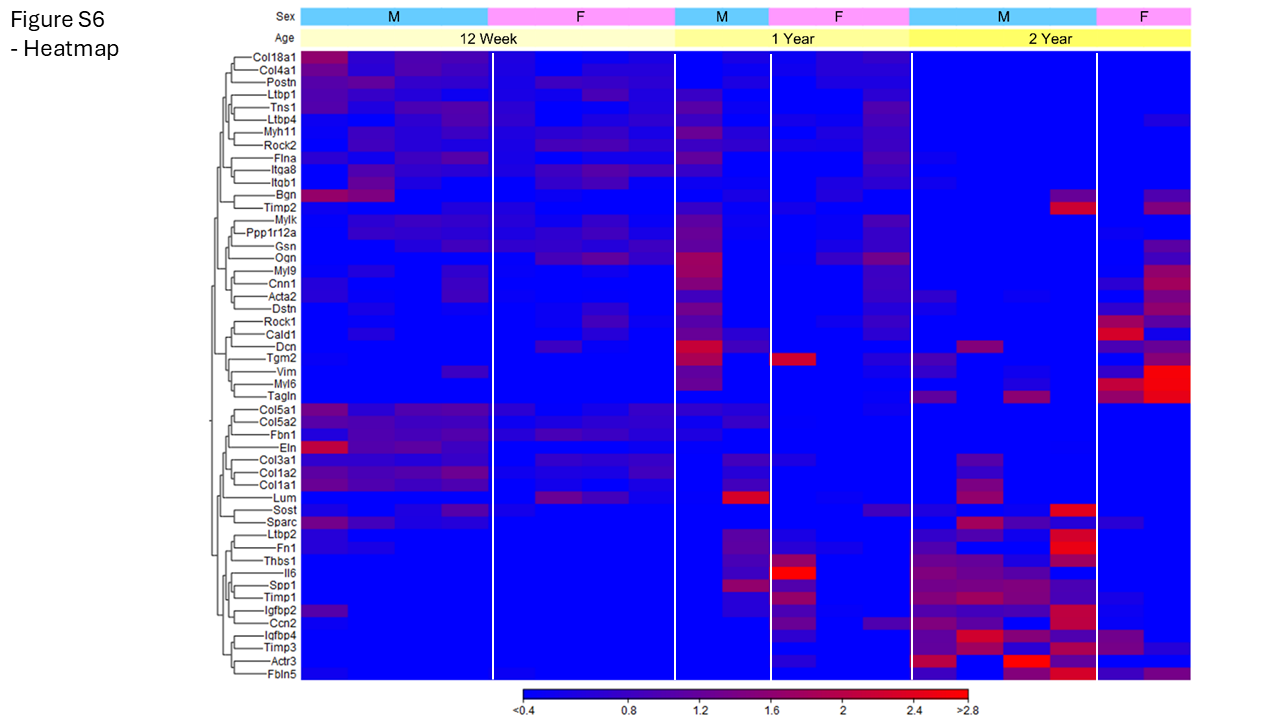

**Figure S6**. Heatmap for male (M) and female (F) Marfan syndrome aortas at all three ages for select transcripts across the three ages.

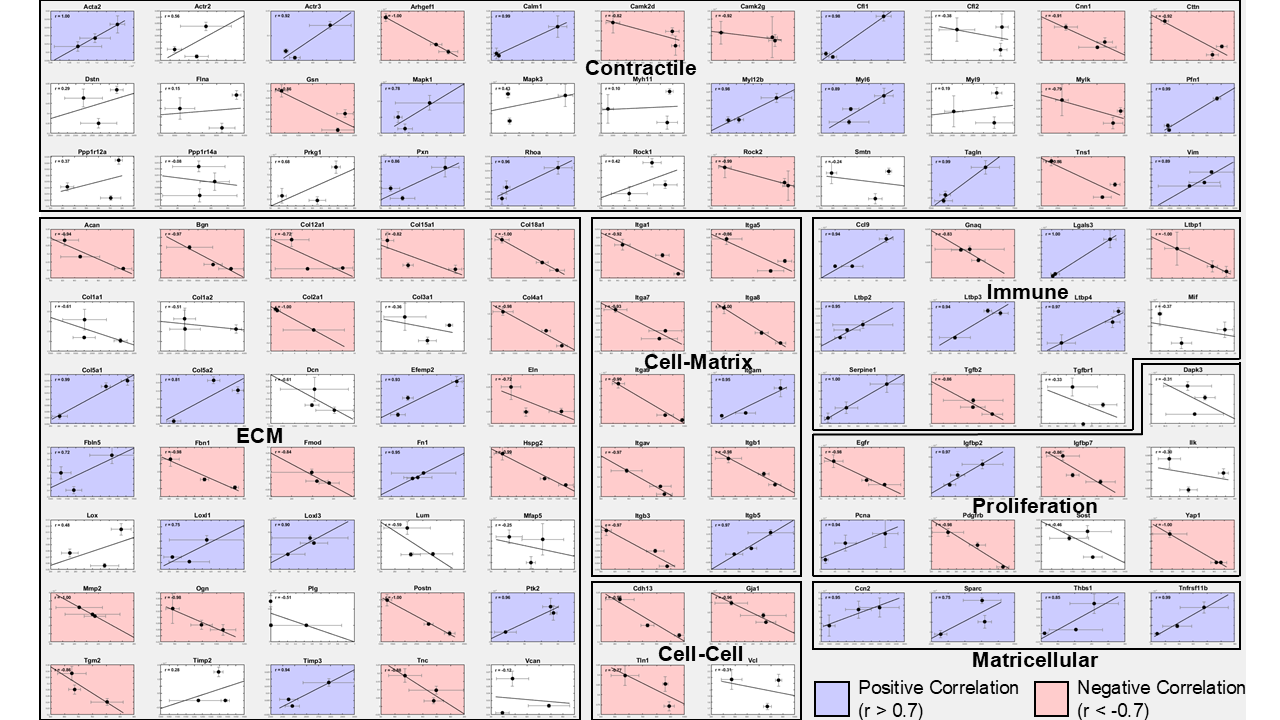

**Figure S7**. Linear correlations, if present, between proteomic and transcriptomic findings across the three ages for MFS aortas. Of the 106 transcripts/proteins shown here, 33% correlated positively (blue), 43% negatively (pink), and 26% not at all (white).

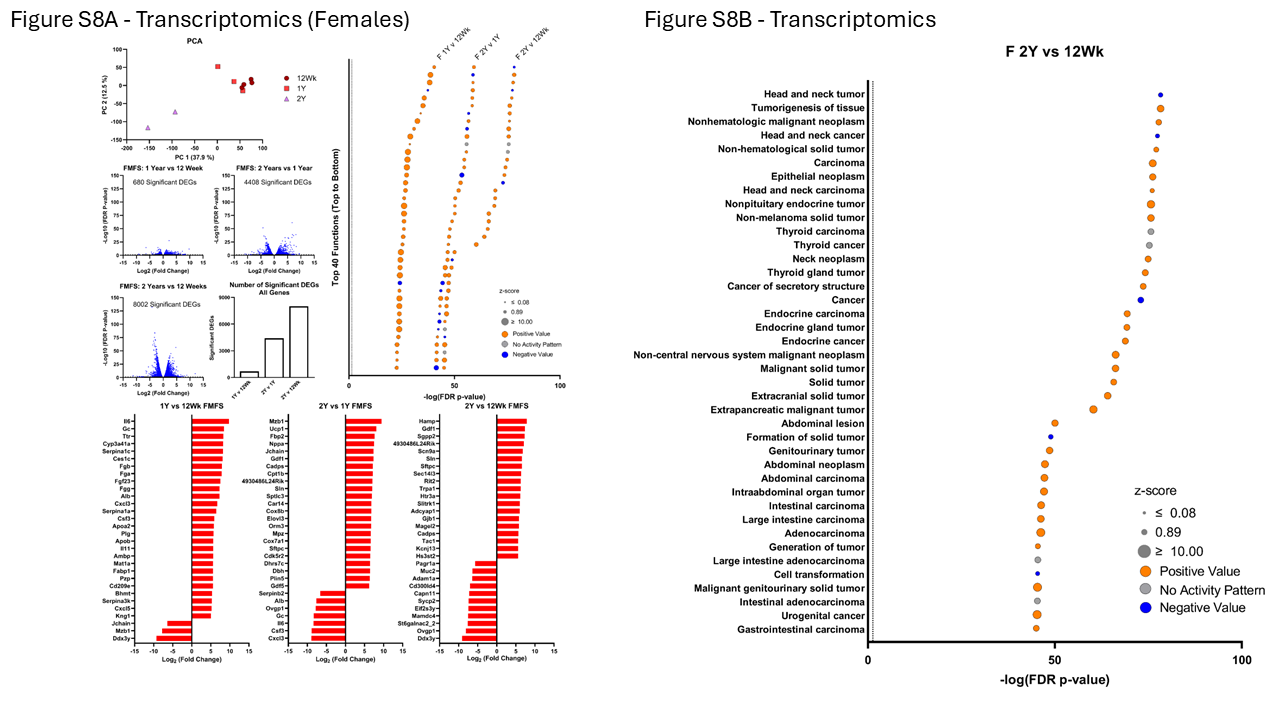

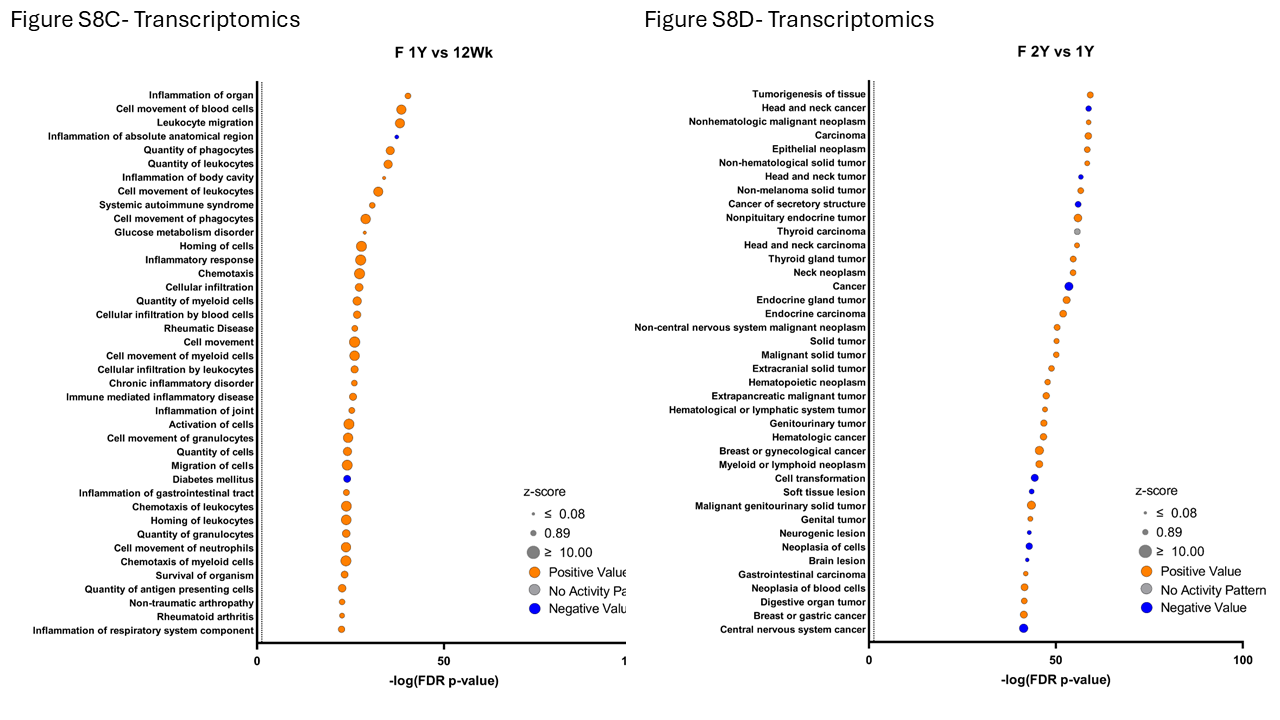

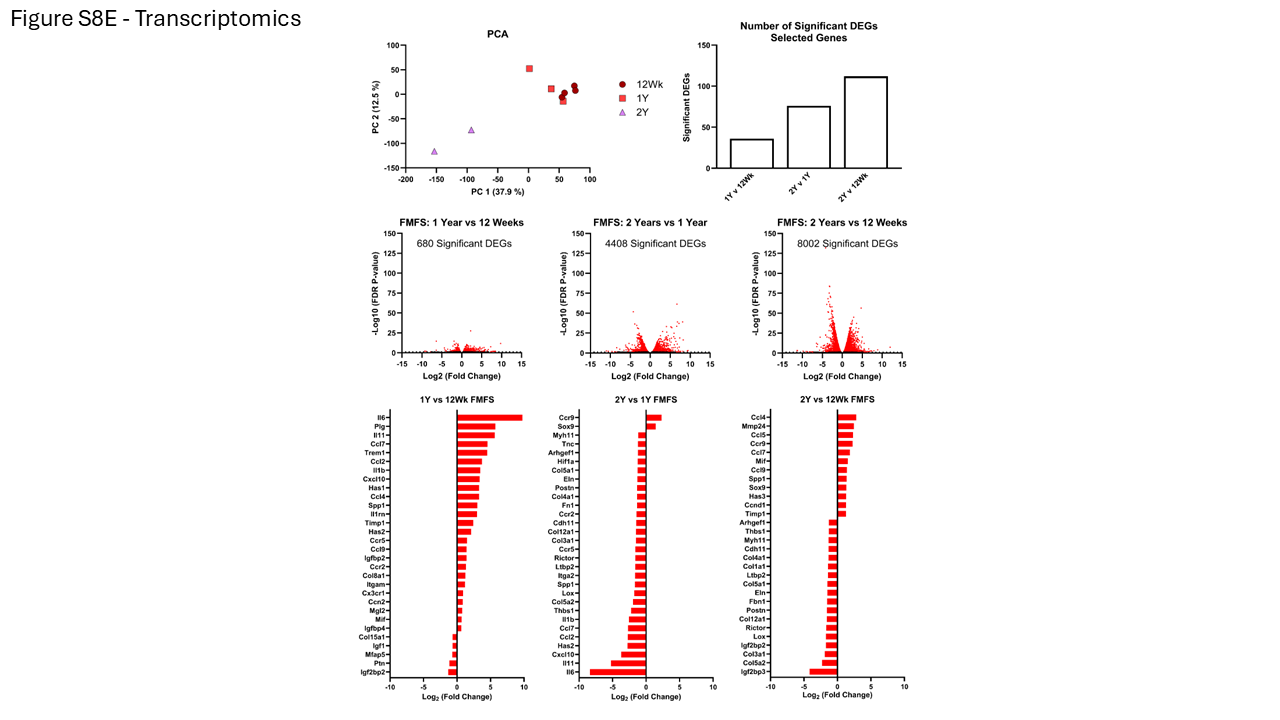

**Figure S8**. Differential gene expression (DEG) across all three ages of female MFS aortas. A. Principal component analysis, volcano plots, and log2 fold changes. B. Associated biological processes for 2 years vs. 12 weeks of age. C. Associated biological processes for 1 year vs. 12 weeks of age. D. Associated biological processes for 2 years vs. 1 year of age. E. Similar to panel A except for select transcripts.

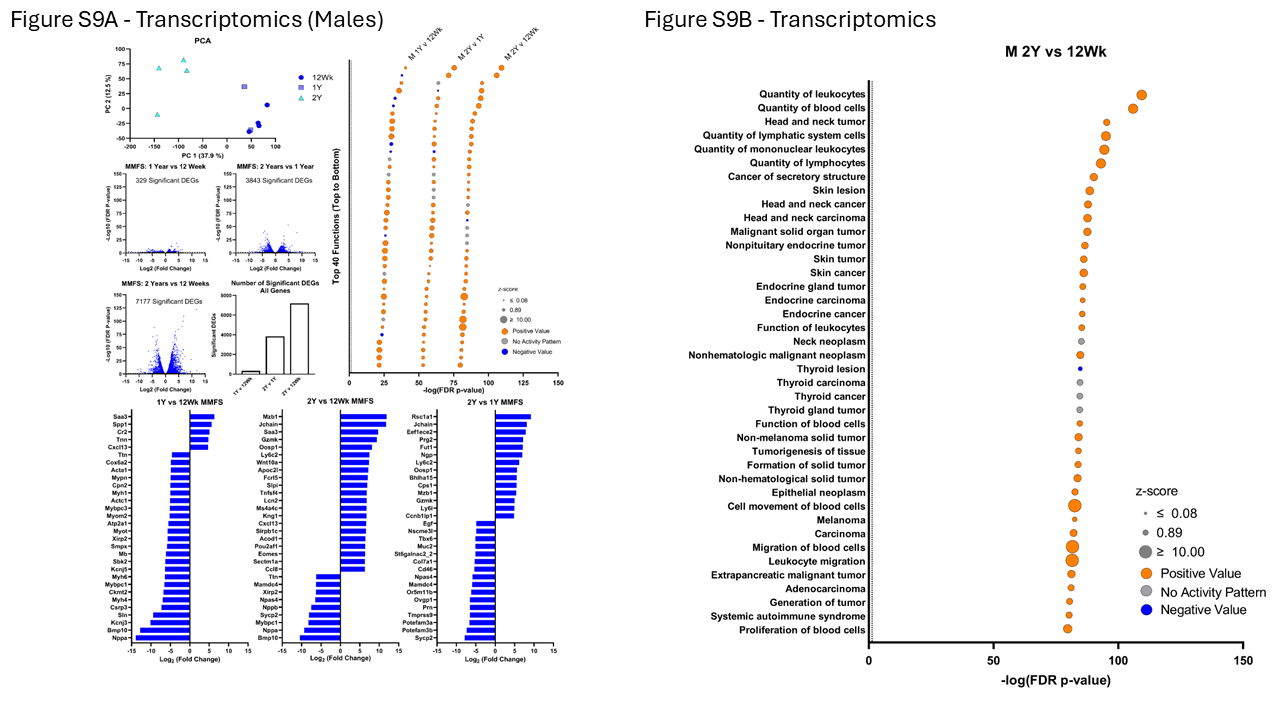

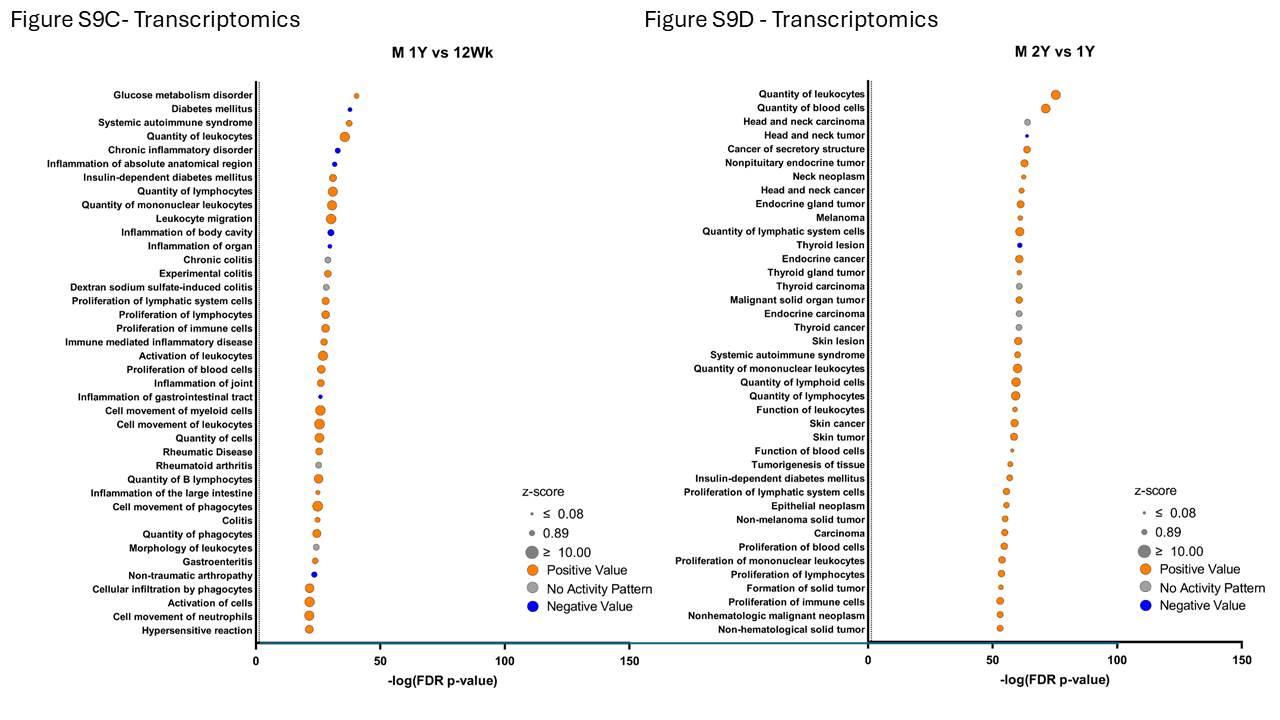

**Figure S9**. Differential gene expression (DEG) across all three ages of male MFS aortas. A. Principal component analysis, volcano plots, and log2 fold changes. B. Associated biological processes for 2 years vs. 12 weeks of age. C. Associated biological processes for 1 year vs. 12 weeks of age. D. Associated biological processes for 2 years vs. 1 year of age. See Figure 5 in the main text for a similar plot for select transcripts.

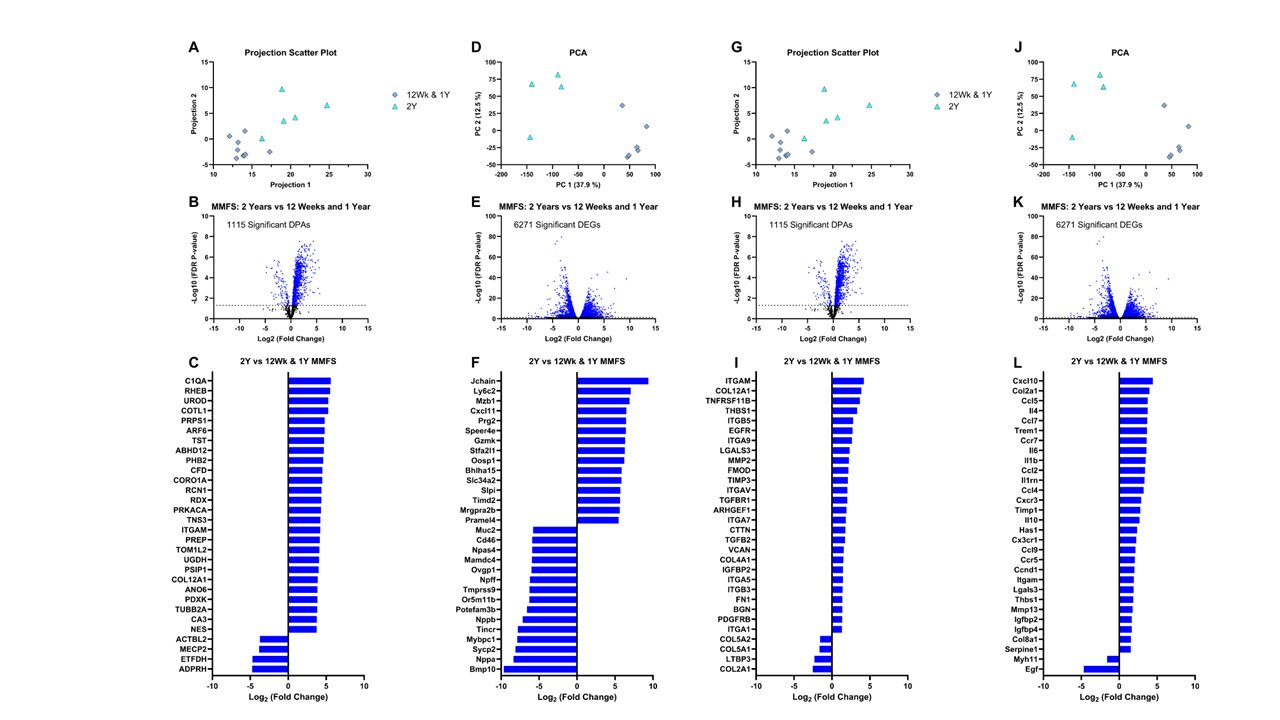

**Figure S10**. Similar to Figures S4 and S9 except for proteomics (left per pairing) and transcriptomics (right per pairing), both in general (left pairs) and for 30 select proteins / transcripts (right pairs) for male MFS mice between two groups: combined 12 weeks + 1 year versus 2 years.
